# DAXX adds a *de novo* H3.3K9me3 deposition pathway to the histone chaperone network

**DOI:** 10.1101/2022.09.20.508668

**Authors:** Massimo Carraro, Ivo A. Hendriks, Colin M. Hammond, Victor Solis, Moritz Völker-Albert, Jonas D. Elsborg, Melanie B. Weisser, Christos Spanos, Guillermo Montoya, Juri Rappsilber, Axel Imhof, Michael L. Nielsen, Anja Groth

**Affiliations:** Novo Nordisk Center for Protein Research (CPR), Faculty of Health Sciences, University of Copenhagen, Copenhagen, Denmark; Biotech Research and Innovation Centre (BRIC), Faculty of Health Sciences, University of Copenhagen, Copenhagen, Denmark; EpiQMAx GmbH, Planegg, Germany; Wellcome Centre for Cell Biology, University of Edinburgh, Edinburgh, UK; Technische Universität Berlin, Chair of Bioanalytics, Berlin, Germany; Biomedical Center - Molecular Biology, LMU Munich

**Author notes:** These authors contributed equally to this work (Massimo Carraro, Ivo A. Hendriks, Colin M. Hammond). Correspondence and requests for materials should be addressed to Anja Groth, Michael Lund Nielsen and Colin Hammond. Lead contact: Anja Groth.

## Abstract

A multitude of histone chaperones are required to protect histones after their biosynthesis until DNA deposition. They cooperate through the formation of co-chaperone complexes, but the crosstalk between nucleosome assembly pathways remains enigmatic. Using explorative interactomics approaches, we characterize the organization of the histone H3–H4 chaperones network and define the interplay between histone chaperone systems. We identify and validate several novel histone dependent complexes and predict the structure of the ASF1 and SPT2 co-chaperone complex, expanding the role of ASF1 in histone dynamics. We show that DAXX acts separately from the rest of the network, recruiting heterochromatin factors and promoting lysine 9 tri-methylation of new histone H3.3 prior to deposition onto DNA. With its functionality, DAXX provides a molecular mechanism for *de novo* heterochromatin assembly. Collectively, our findings provide a new framework for understanding how cells orchestrate histone supply and comply with chromatin dynamics throughout the cell cycle.

## Introduction

In eukaryotic cells, genomic DNA is packaged with histone proteins into chromatin which regulates genome function and stability. The basic repeating unit of chromatin is the nucleosome, formed by 147 base pairs of DNA wrapped around an octameric complex of histones H3, H4, H2A and H2B (Luger et al., 1997). Nucleosomes are modified through histone post-translational modifications (PTMs) and the substitution of core histones with histone variants. This histone-based epigenetic information drives chromatin functionality, regulating gene expression, silencing repetitive elements, and instructing DNA damage response pathways (Lai and Pugh, 2017; Nicetto and Zaret, 2019). To allow the passage of the molecular machines that transcribe, replicate, or repair the DNA template, nucleosomes are disassembled and reassembled, and this is complemented with new histone deposition pathways that maintain nucleosome density (Grover et al., 2018; Hammond et al., 2017; Pardal et al., 2019). This is especially important during DNA replication where deposition of new histones is required to maintain nucleosome density on new daughter DNA strands (Stewart-Morgan et al., 2020).

Histone supply and chromatin dynamics are supported by a structurally diverse set of proteins called histone chaperones (Grover *et al.*, 2018; Hammond *et al.*, 2017; Pardal *et al.*, 2019). Histone chaperones shield the interactions of histones with DNA/RNA in a manner that only proper nucleosome contacts can out-compete (Andrews et al., 2010), thereby promoting the ordered assembly and disassembly of nucleosomes. Histone chaperones often collaborate, forming histone dependent co-chaperone complexes (Hammond *et al.*, 2017). In these complexes, multiple chaperones simultaneously associate with the same histone-fold dimer or tetramer providing a more complete shield around the histone substrate (Hammond et al., 2016; Huang et al., 2015; Ricketts et al., 2015; Saredi et al., 2016), and potentially promoting nucleosome assembly (Huang *et al.*, 2015). Histone chaperones can also combine through direct histone independent interactions to provide multivalency to their chaperoning functionality or to mediate histone handover events. Similarly, histone chaperone functionality integrates with chromatin remodelers, histone modifying enzymes, heat shock molecular chaperones, and DNA/RNA polymerases and helicases (Hammond et al., 2021; Hammond *et al.*, 2017; Loyola et al., 2009), influencing histone deposition and chromatin states.

In mammals, the incorporation of the canonical histone H3 (H3.1 and H3.2) and its replacement variants (H3.3 and CENPA) have profound effects on chromatin organization (Martire and Banaszynski, 2020), and each variant associates with partially distinctive chaperone systems. After translation, newly synthesized canonical histones H3.1/2 and variant H3.3 are co-folded with histone H4 by the histone chaperone DNAJC9 (Hammond *et al.*, 2021). Folded H3–H4 dimers are then handled by histone chaperone NASP, which protects a soluble pool of histones from chaperone-mediated autophagy (Bao et al., 2022; Cook et al., 2011; Hormazabal et al., 2022). The somatic isoform of NASP (sNASP) also functions in complex with histone chaperones RbAp46 (RBBP7) and the histone acetyltransferase HAT1 (Campos et al., 2010). The HAT-1 complex promotes histone H4 K5/K12 acetylation of H3–H4 dimers which associate with ASF1 for nuclear import via Importin-4/IPO4 (Grover *et al.*, 2018; Hammond *et al.*, 2017). ASF1 coordinates *de novo* H3–H4 supply to the CAF-1 and HIRA complexes (Grover *et al.*, 2018; Hammond *et al.*, 2017), which are responsible for replication- and transcription-coupled deposition of H3.1/2–H4 and H3.3–H4 respectively (Tagami et al., 2004).

During histone supply, ASF1 also forms co-chaperone interactions with both MCM2 and TONSL (Groth et al., 2007; Huang *et al.*, 2015; Jasencakova et al., 2010; Saredi *et al.*, 2016). MCM2 is part of the CMG helicase complex (CDC4, MCM2-7 and GINS) and functions as a histone recycling factor during DNA replication (Gan et al., 2018; Petryk et al., 2018), while stabilizing soluble H3–H4 bound by ASF1 (Huang *et al.*, 2015). TONSL binds newly synthesized histones through recognition of histone H4 unmethylated at lysine 20 (Saredi *et al.*, 2016), a mark of post-replicative chromatin that promotes DNA damage repair via homologous recombination (Nakamura et al., 2019; Saredi *et al.*, 2016). Whilst ASF1, MCM2 and TONSL are co-chaperone partners (Huang *et al.*, 2015; Saredi *et al.*, 2016), it is not entirely clear whether TONSL contributes to both H3.1/2 and H3.3–H4 supply pathways. To add to the complexity, ASF1 has two isoforms, ASF1a and ASF1b, that have partially overlapping functions in histone supply (Groth et al., 2005), but may cooperate differently with downstream histone chaperones.

At sites of constitutive heterochromatin H3.3–H4 dimers are deposited by the histone chaperone DAXX (Drané et al., 2010; Goldberg et al., 2010; Lewis et al., 2010). DAXX-mediated H3.3–H4 deposition is essential for maintaining silencing of repetitive DNA elements including telomeres, viral genomes, retrotransposons and imprinted regions (Elsässer et al., 2015; He et al., 2015; Tsai and Cullen, 2020; Voon et al., 2015), and also plays a role in protecting stalled replication forks (Teng et al., 2021). Despite the importance of this deposition pathway, the stage of histone supply when DAXX associates with histones remains elusive (Hammond *et al.*, 2017). DAXX interacts with the chromatin remodeler ATRX (Dyer et al., 2017), the histone methyltransferases SUV39H1 (He *et al.*, 2015) and SETDB1/SETB1/ESET, and the SETDB1-linked co-repressor protein TRIM28/TIF1B/KAP1 (Elsässer *et al.*, 2015; Hoelper et al., 2017) during the establishment of heterochromatic silencing. Additionally, DAXX-mediated transcriptional silencing requires DAXX localization to Promyelocytic Leukemia (PML) nuclear bodies in a SUMOylation dependent manner (Corpet et al., 2020). DAXX deposition of H3.3-H4 is required for maintenance of H3K9me3 (Elsässer *et al.*, 2015; Groh et al., 2021; He *et al.*, 2015; Sadic et al., 2015), a PTM linked to transcriptional silencing (Padeken et al., 2022). Why H3.3 deposition is required for H3K9me3 enrichment and heterochromatin silencing remains unclear, especially since H3.3–H4 is also deposited at sites of active transcription by HIRA (Goldberg *et al.*, 2010). CENPA–H4 dimers are also handled in a variant-specific manner by the histone chaperone HJURP (Dunleavy et al., 2009; Foltz et al., 2009), which promotes their deposition at centromeric chromatin through the MIS18 complex (Barnhart et al., 2011; Fujita et al., 2007). The histone chaperones RBBP4/7 and NPM also collaborate during CENPA–H4 supply (Dunleavy *et al.*, 2009; Foltz *et al.*, 2009), but their interplay with HJURP is not defined.

The cooperative nature of histone chaperones in histone metabolism suggests the existence of an interconnected chaperone network (Hammond *et al.*, 2017). However, the organization of the histone chaperone network and the crosstalk between H3 variant supply pathways has not been systematically studied. Here, we define the topology of the histone chaperone network surrounding key nodes in the histone H3 variant supply chains, providing a rich resource to broaden our understanding of new and existing players in histone chaperone biology. We delineate the crosstalk between different H3–H4 chaperones systems revealing that DAXX operates as a largely independent arm of the histone chaperone network. Through interrogation of DAXX functionality, we demonstrate a route for the delivery of newly synthesised histones H3.3 modified with K9me3 to chromatin, unveiling a molecular mechanism for *de novo* heterochromatin assembly.

## Results

### Charting the histone chaperone network

To understand the connectivity within the histone chaperone network and identify new co-chaperone relationships, we profiled ASF1a/b, sNASP, HJURP and DAXX histone-dependent and -independent interactomes. Together this panel of histone chaperones, allowed us to monitor the pathways of replication dependent and independent nucleosome assembly, soluble histone homeostasis, centromere assembly and heterochromatin maintenance (Grover *et al.*, 2018). To this end, we compared the interactomes of conditionally expressed wild type (WT) histone chaperones to their corresponding histone binding mutant (HBM), and a negative control in triple SILAC IP-MS experiments (Stable Isotope Labeling with Amino acids in Cell culture, ImmunoPrecipitation coupled to Mass Spectrometry) (**Figure 1A)**. Based on previous studies, we identified point mutations that disrupt histone binding for these histone chaperones to use in our proteomic comparisons (**Figure S1A**) (Bowman et al., 2016; Elsässer et al., 2012; English et al., 2006; Hu et al., 2011). We recently identified DNAJC9 as a new player in the histone chaperone network using a similar experimental strategy (Hammond *et al.*, 2021).

**Figure 1.**
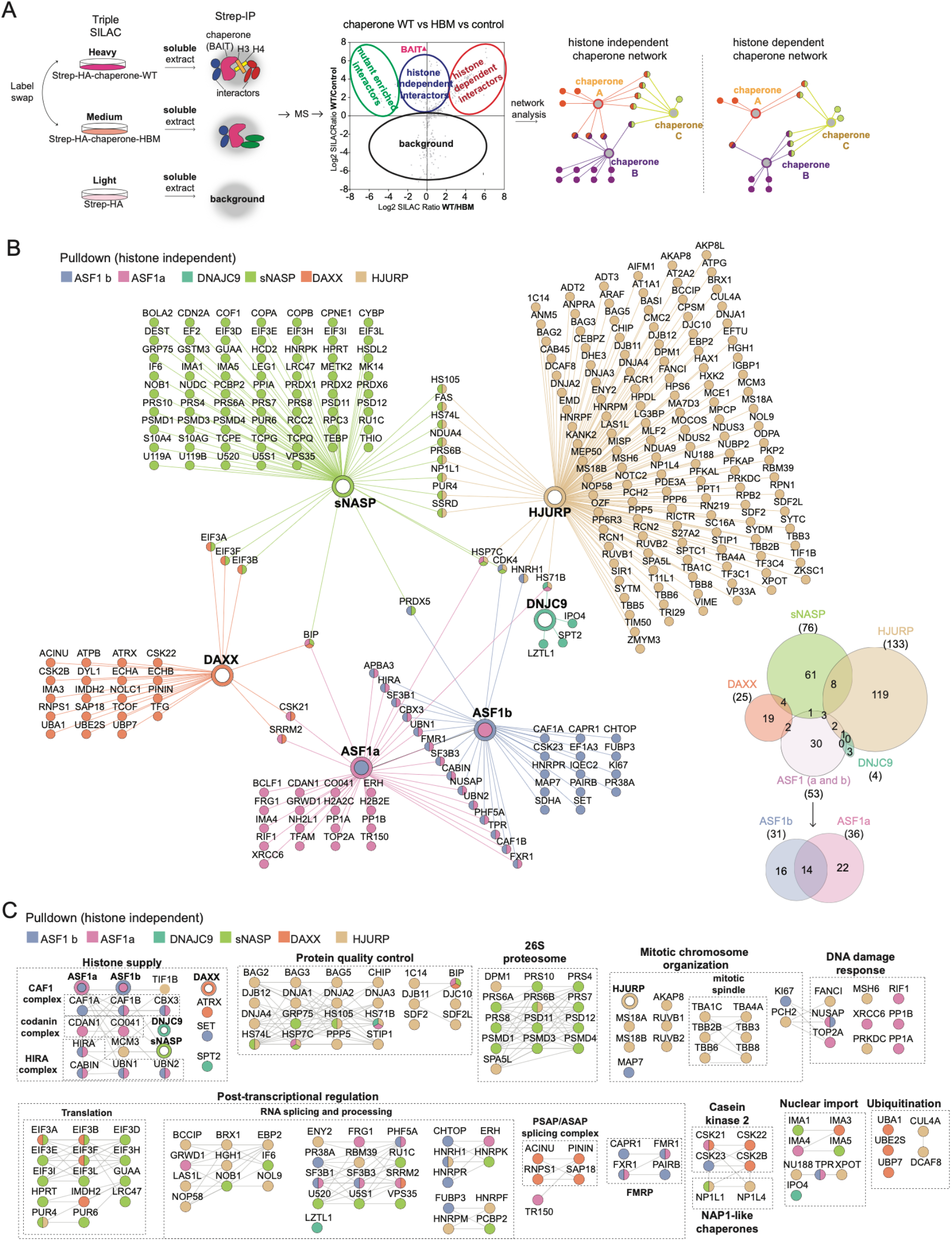
Histone chaperones have diverse histone independent functions. (A) The triple SILAC IP-MS strategy for mapping the histone chaperone network. (B) (Left) Network analysis of the histone independent interactors identified for ASF1a, ASF1b, NASP, DAXX, HJURP and DNAJC9. (Right) Venn diagram showing the overlaps between histone independent interactomes. Proteins nodes, edges and Venn categories are colored based on their identifications in the different pulldowns. (C) Clustering of histone independent interactors using functional associations annotated in the STRING database and the MCL algorithm. Protein-protein functional associations are shown with a black line, according to the string database. (B-C) Proteins are referred to by human UniProt protein identification code. Data generated from n=4 biological replicates. See also **Table S1** and **Figure S2A**.

We compared our datasets to findings from the literature, confirming the robust nature of our experimental approach (**Figure S1B-F, Table S1**). For instance, we were able to confirm ASF1a/b histone dependent interactions with MCM2 (Huang *et al.*, 2015), TONSL (Saredi *et al.*, 2016), NASP and the HAT-1 complex (HAT1 and RBBP7), and IPO4 (Campos *et al.*, 2010; Jasencakova *et al.*, 2010) (**Figure S1B-C**). sNASP interactomes corroborated the histone dependent association with ASF1a/b and other HAT-1 complex members (Bowman et al., 2017; Campos *et al.*, 2010; Jasencakova *et al.*, 2010) (**Figure S1D**). Furthermore, ASF1a/b also co-purified CAF1B (CAF-1 p60) and the HIRA complex (HIRA, UBN1/2, CABIN) independent of histone binding (**Figure S1B-C**), in line with expectations (Tang et al., 2006). Finally, our pulldowns confirmed the direct histone independent interaction of DAXX and ATRX (**Figure S1F**) (Hoelper *et al.*, 2017; Lewis *et al.*, 2010).

We next performed a network analysis of histone independent interactors identified in our histone chaperone interactomes (**Figure 1B, Table S1**), including the published triple SILAC IP-MS datasets for DNAJC9 (Hammond *et al.*, 2021). Strikingly, most of the histone chaperones analysed had a largely unique histone independent interactome, with only a few proteins interacting with multiple chaperones. To explore the biological themes encoded in these interactomes, we generated a clustered interaction network that integrates the functional associations between interactors from the STRING database. We then mapped these clusters to known protein complexes and pathways highlighting both common and distinct biological pathways across the different histone chaperone interactors. (**Figure 1C, Figure S2A**). This revealed an enrichment of proteins involved in post-transcriptional regulation and RNA processing in several of the interactomes. For instance, DAXX directly associates with PSAP and ASAP splicing complexes (SAP18, PININ, RNPS1, ACINU) (Murachelli et al., 2012), NASP associates with several pre-mRNA binding proteins (RUC1, U520, U5S1, and HNRPK), and ASF1a/b link to Fragile X syndrome RNA binding proteins (FXR1, FMR1, CAPR1, PAIRB) (Hagerman et al., 2017). This underscores the integration of histone chaperones with post-transcriptional regulation as an emerging theme in histone chaperone biology (Kim et al., 2018; Park et al., 2018). The histone independent enrichment of heat shock molecular chaperones was another common theme shared across datasets, strengthening the functional link between these complementary chaperone systems (Hammond *et al.*, 2021). Meanwhile, nuclear import proteins also featured prominently in this analysis. sNASP interacted with the Importin proteins IMA1/KPNA2 and IMA5/KPNA1, the latter was recently implicated in the nuclear import of monomeric histones (Pardal and Bowman, 2021), which can also be bound in the nucleus by sNASP (Apta-Smith et al., 2018). DNAJC9 formed histone independent interactions with Importin-4 (IPO4) (Hammond *et al.*, 2021), while ASF1 and DAXX associated with the importins IMP4/KPNA3 and IMA3/KPNA4 respectively, potentially providing alternative pathways for the nuclear import of histone H3–H4 dimers.

In contrast, several unique biological processes were enriched by an individual histone chaperone or a select subset of those profiled. Several subunits of the 26S proteosome (PRS4, PRS6A, PRS6B, PRS7, PRS8, PRS10, PRS11, PRS12; PSMD1, PSMD3, PSMD4)(Tanaka, 2009) were exclusively enriched with sNASP, placing NASP in proximity of both the proteasomal and autophagy-mediated (Cook *et al.*, 2011) degradation machineries. Proteins involved in mitotic chromosome segregation (MIS18A, MIS18B, TRIP13, RUVB1, RUVB2, AKAP8) (Collas et al., 1999; Magalska et al., 2014; Müller and Almouzni, 2014; Yost et al., 2017) and the mitotic spindle assembly (TBA1C, TBA4A, TBB2B, TBB3, TBB6, TBB8) (Prosser and Pelletier, 2017) specifically associate with HJURP. These histone independent interactions are in line with the histone binding domain of HJURP being dispensable for its recruitment to the MIS18 complex (MIS18A and MIS18B)(Pan et al., 2019), the complex required for HJURP centromere localization (Barnhart *et al.*, 2011; Fujita *et al.*, 2007). Finally, factors involved in DNA repair pathways were purified with either HJURP (MSH6, FANCI, PRKDC) or ASF1a (XRCC6, TOP2A, RIF1 and PP1A/B), consistent with the involvement of these two chaperones in DNA repair pathways (Lee et al., 2017; Yilmaz et al., 2021). To cross-validate the isoform specificity of ASF1 interactors, we directly compared ASF1a and ASF1b interactomes (**Figure S2B**). Corroborating previous findings, ASF1b had a preference for the CAF-1 complex (Abascal et al., 2013) and ASF1a associates specifically with RIF1 in a histone-independent manner (Tang et al., 2022), and suggests RIF1 could recruits ASF1a to specific genomic position for histones deposition. Meanwhile, the HIRA complex associates with both ASF1 isoforms, consistent with our previous experiment (**Figure 1B**).

### Histone chaperone cooperation is built through histone dependent interactions

Next, we compared the histone dependent interactors identified in our chaperone pulldowns, incorporating published datasets for DNAJC9, MCM2 and TONSL (Hammond *et al.*, 2021). This analysis revealed a more substantial overlap in interactors (**Figure 2A**), demonstrating the robust crosstalk between histone supply pathways. ASF1a and ASF1b shared most of the histone dependent interactors, in line with the similar roles of the two isoforms in nucleosome assembly pathways (Groth *et al.*, 2005). Many of these ASF1 interactors were also histone-dependent interactors of DNAJC9, corroborating the idea that DNAJC9 operates in parallel to ASF1 in histone supply (Hammond *et al.*, 2021). Despite the high degree of overlap between chaperones in the histone dependent network we only observed two interactions linking DAXX to the rest of the network, NP1L4/NAP1L4 and C1QBP, shared with ASF1a and ASF1b. As NAP1L4 and C1QBP are known histones chaperones that bind H3–H4 (Lin et al., 2021; Okuwaki et al., 2010), they represent a possible link between DAXX and ASF1 nucleosome assembly pathways.

**Figure 2.**
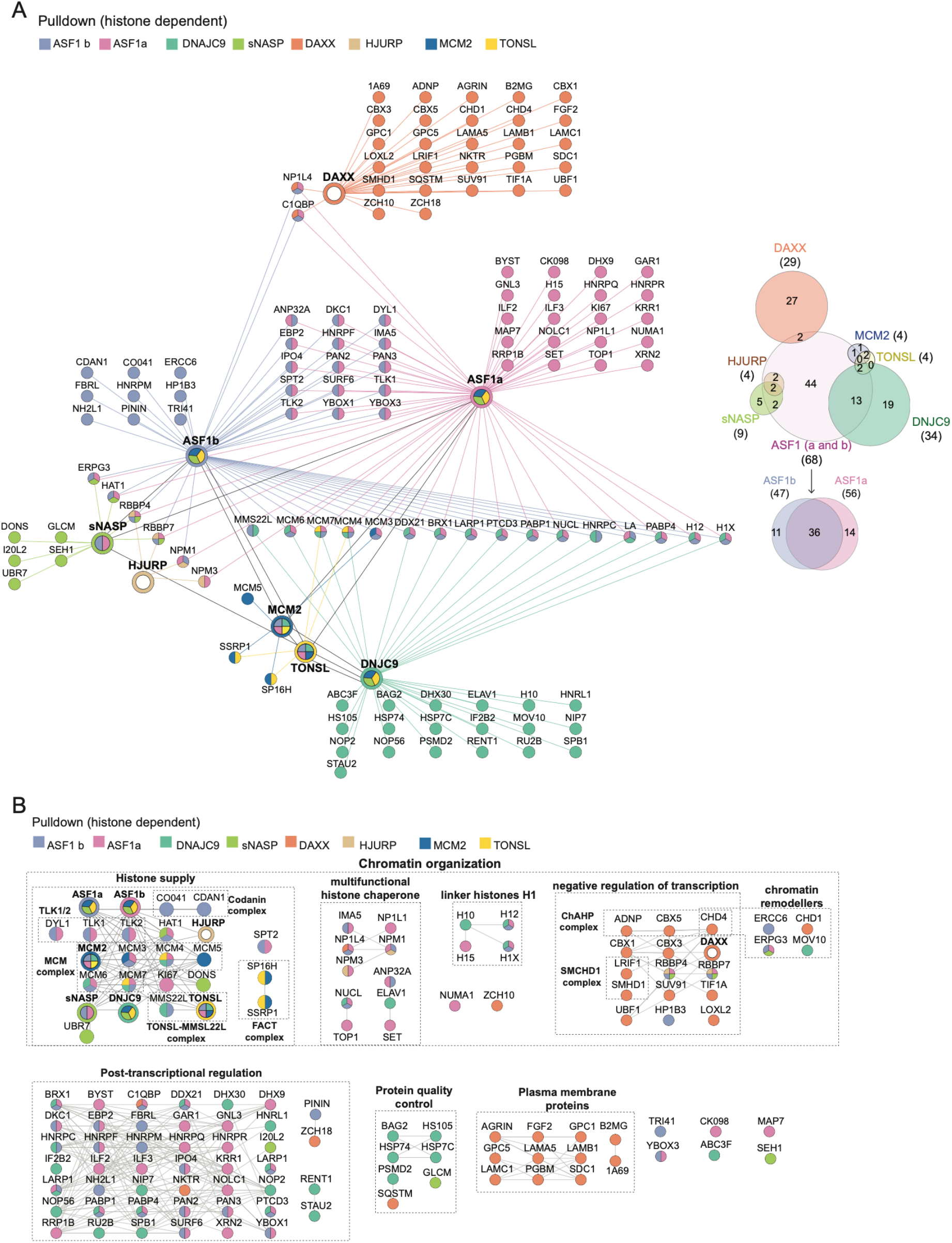
Histone chaperones collaborate through their histone dependent associations. (A) (Left) Network analysis of the histone dependent interactors identified for ASF1a, ASF1b, NASP, DAXX, HJURP, DNAJC9, MCM2 and TONSL. (Right) Venn diagram showing the overlaps between histone dependent interactomes. Proteins nodes, edges and Venn categories are colored based on their identifications in the different pulldowns. (B) Clustering of histone independent interactors using functional associations annotated in the STRING database and the MCL algorithm. Protein-protein functional associations are shown with a black line, according to the string database. (A-B) Proteins are referred to by human UniProt protein identification code. Data generated from n=4 biological replicates. See also **Table S1**.

We grouped the histone dependent proteins into known pathways and complexes, defining the common and unique features of the histone delivery systems (**Figure 2B**). Again, an enrichment of factors involved in post-transcriptional was observed, but this time in a histone dependent manner, demonstrating that histone chaperone functionality is linked to the RNA processing machinery. DNAJC9, sNASP and DAXX were associated in a histone-dependent manner with factors involved in protein quality control. DNAJC9 was the only chaperone to contact heat shock molecular chaperones in a histone-dependent manner (HSP74, HSP7C, HS105 and BAG2), supporting its unique role in recruiting these molecular chaperones to soluble histones (Hammond *et al.*, 2021). sNASP and DAXX purified with factors involved in autophagy-mediated protein degradation (GLCM and SQSTM, respectively). sNASP is known to protect soluble histones H3 and H4 from chaperone-mediated autophagy (Cook *et al.*, 2011; Hormazabal *et al.*, 2022), thus the histone dependent association with the Lysosomal acid glucosylceramidase GLCM (Boer et al., 2020) is surprising. However, along with the histone-independent association of sNASP and the proteasome (**Figure 1C**), this suggests that sNASP may coordinate a protein folding versus degradation decision point for soluble histones.

Meanwhile, DAXX formed histone dependent interactions with a set of enigmatic proteins, including the proteoglycans (GPC1 and GPC5) (Filmus, 2001), and major histocompatibility complex I molecules (1A69/HLAA and beta-2-microglobulin/B2MG)(Pishesha et al., 2022), and the cargo receptor for selective autophagy SQSTM/p62 (Sánchez-Martín et al., 2019). These interactors suggest a role of DAXX in the membrane localization or extracellular secretion of histones (Chen et al., 2014; Silvestre-Roig et al., 2019), potentially utilized in the innate immune response to silence incoming viral genomes. Indeed, several lines of evidence support a role for DAXX in silencing viral genomes and DAXX is targeted by viral proteins to overcome host immunity (Schreiner and Wodrich, 2013). Otherwise, DAXX–H3–H4 soluble complexes were predominantly associated with factors involved in negative regulation of transcription (CBX5/HP1α, CBX1/HP1β, CBX3/HP1γ, TIF1A/TRIM24, ADNP, SUV91/SUV39H1, SMHD1/SMCHD1, LRIF1, LOXL2)(Allshire and Madhani, 2018). This argues that DAXX specifically recruits factors involved in heterochromatin organization to its H3.3–H4 cargo, prior to their deposition on DNA.

### Novel histone dependent interactions and histone co-chaperone complexes

Our network analysis also enabled the identification of several uncharacterized histone co-chaperone complexes (**Figure 3A**). For example, ASF1a associated with SPT2, AN32A, SET, NP1L1/NAP1L1, NP1L4/NAP1L4, NPM1/NPM, NPM3 and NUCL, of which only NP1L1 and NPM3 were not observed with ASF1b. The histone dependent association of ASF1 isoforms with Nap1-Like proteins (NAP1L1 and NAP1L4), parallels the abilities of both yeast Nap1 and Vps75 to form co-chaperone complexes with Asf1–H3–H4 (Hammond *et al.*, 2016), attesting that these Nap1-Like proteins also bind H3–H4 dimers in mammals. NAP1L4 and NPM1/3 also formed co-chaperone complexes with DAXX and HJURP, respectively, positioning these multifunctional chaperones at the intersection between branches of the H3–H4 network. Considering the histone variant specificities of HJURP and ASF1 towards CENPA–H4 and H3.1/2/3–H4 respectively, their shared histone-dependent associations with NPM1/3 could be bridged by histone H4 or conserved regions of H3 and CENP-A. In line with our results, NPM1 has also been shown to associate with both non-nucleosomal H3–H4 and CENPA–H4 (Dunleavy *et al.*, 2009; Foltz *et al.*, 2009). Finally, sNASP interacted with the novel histone reader UBR7 via histones, demonstrating the histone dependence of this recently reported interaction (Hogan et al., 2021).

**Figure 3.**
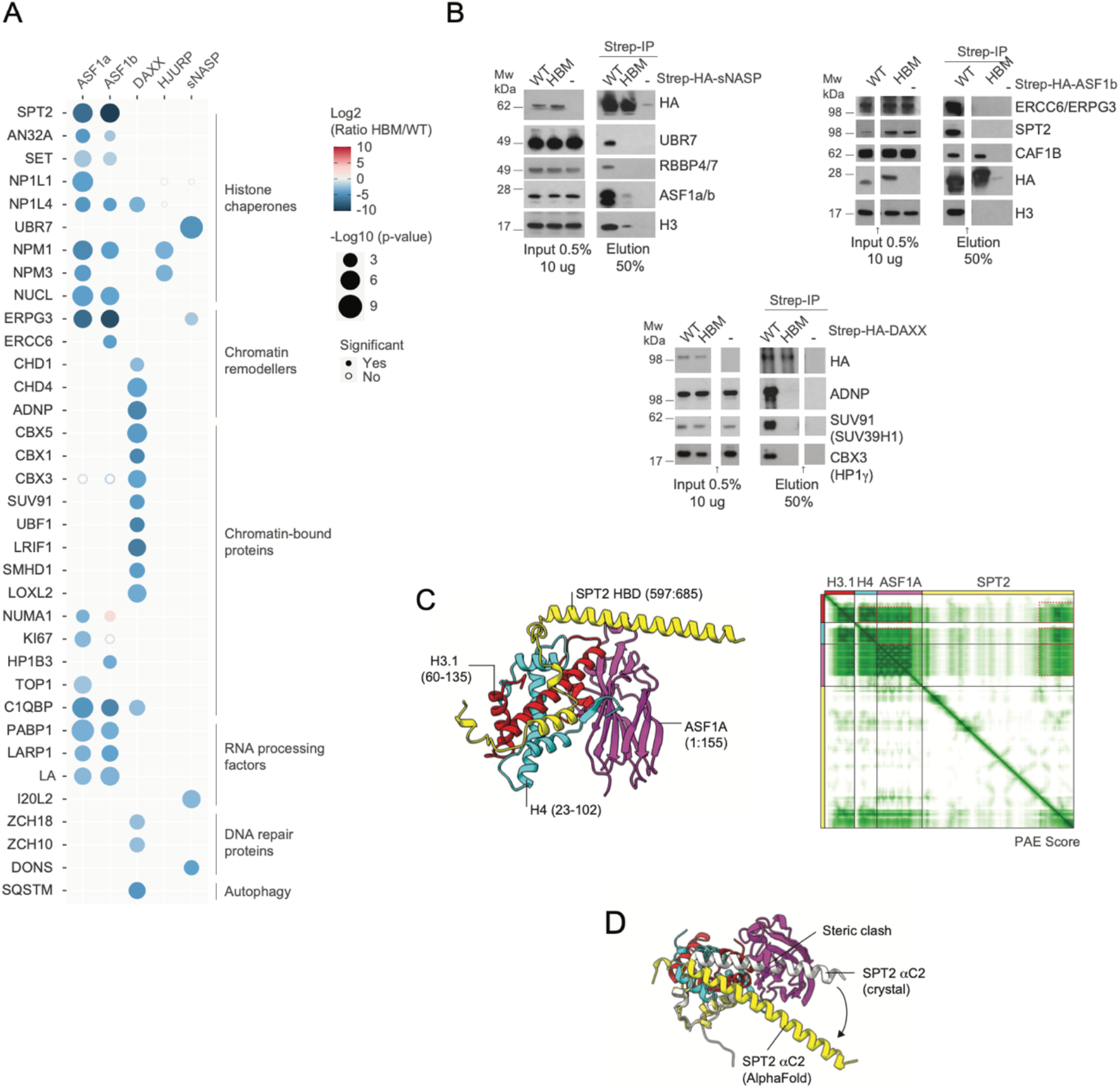
Validation of novel histone dependent interactors. (A) Bubble plot showing the histone dependence of novel histone dependent complexes across triple SILAC IP-MS interactomes. Proteins are referred to by human UniProt protein identification code. Data generated from n=4 biological replicates. See also **Table S1**. (B) Pull-downs of Strep-HA-tagged histone chaperones WT or HBM compared with control purifications (−) from soluble cell extracts probed by Western blot. Representative of n = 2 biological replicates. Arrow indicates lanes containing unrelevant samples have been removed. See also **Figure S3A-C**. (C) (left) ‘Local’ AlphaFold prediction of SPT2 (yellow) and ASF1A (magenta) histone binding domains bound to H3.1–H4 (red and cyan) depicting the high confidence regions of the full-length AlphaFold prediction. (right) Predicted Alignment Error (PAE) plot showing confidence of residue contacts in the full-length SPT2–H3.1–H4–ASF1A ‘global’ AlphaFold prediction. Red dashed lines indicate high confidence interactions between protein chains in the ‘local’ prediction shown (left). (D) Alignment of local AlphaFold prediction of SPT2–H3.1–H4–ASF1A (colored as per panel C) to the crystal structure of SPT2– (H3.2–H4)_2_ (white; PDB: 5BS7, with H3.2–H4 omitted for clarity). See also **Figure S3D-E**.

In addition, we identified several histone dependent interactions with chromatin remodelers. DAXX has histone dependent relationships with the chromatin remodelers CHD1 and the ChAHP complex (CHD4, ADNP, CBX1/3). With the ability of DAXX to deposit H3.3–H4 in collaboration with the chromatin remodeler ATRX, these links to other chromatin remodeling enzymes could allow H3.3–H4 deposition by DAXX at other genomic sites. In addition, the chromatin remodeler ERCC6/CSB (and its alternative splicing product, ERPG3) was identified in ASF1a/b and sNASP histone dependent complexes. ERCC6 remodels chromatin during transcription-coupled nucleosome excision repair (Hargreaves and Crabtree, 2011), and our result suggests ASF1 and sNASP assist in this process.

To further the resource potential of our histone chaperone network analysis, we validated the histone dependent interactions of ASF1b with ERCC6/ERPG3 and SPT2, and sNASP interaction with UBR7 by western blotting (**Figure 3B**). Moreover, we confirmed histone-dependent interactions of DAXX with HP1y (CBX3), ADNP and SUV39H1 (SUV91), validating DAXX recruitment of these heterochromatin factors to its histone substrate (**Figure 3B**). Finally, we investigated the molecular bases of ASF1 and SPT2 co-chaperone complex. SPT2 is an H3–H4 histone chaperone involved in transcription-coupled histone dynamics, required for maintaining chromatin organization across gene bodies and preventing spurious transcription events (Chen et al., 2015; Osakabe et al., 2013). Using AlphaFold 2.0 multimer (Evans et al., 2021; Jumper et al., 2021) we predicted the structure of the ASF1a–H3.1–H4– SPT2 co-chaperone complex (**Figure S3A-B**). From this, we identified an interacting region containing the histone binding domains of SPT2, ASF1 and the histone fold of H3.1–H4 which generated high confidence scores in both PAE and pLDDT metrics (**Figure 3C**, **Figure S3A-C**). Alignment of this AlphaFold prediction with the crystal structure of SPT2–(H3.2–H4)_2_ indicated that the SPT2 αC2 helix (**Figure 3D**, **Figure S3D)**, normally associated with the H3– H4 tetramerization interface, is relocated to allow ASF1a to maintain binding mode with H3– H4 (**Figure S3E**) (English *et al.*, 2006; Natsume et al., 2007). This prediction supports the simultaneous binding of ASF1a/b and SPT2 to the same histone substrate, arguing the two histone chaperones could cooperate during transcription-coupled nucleosome assembly/disassembly (Chen *et al.*, 2015; Osakabe *et al.*, 2013).

### DAXX operates as an isolated arm of the histone H3–H4 chaperone network

To further dissect the role of ASF1a, ASF1b, NASP and DAXX in the histone supply chain, we characterized the effect of their depletion on the interactomes of soluble H3.1 and H3.3, using label free quantification (LFQ) coupled MS analysis (**Figure 4A-C**). We focused our analysis on well-known histone H3–H4 chaperones and their associated binding partners, several importins (IPO4, IMA3, IMA4, IMA5), and some more recently implicated histone binding factors (UBR7, C1QBP) (Hogan *et al.*, 2021; Lin *et al.*, 2021).

**Figure 4.**
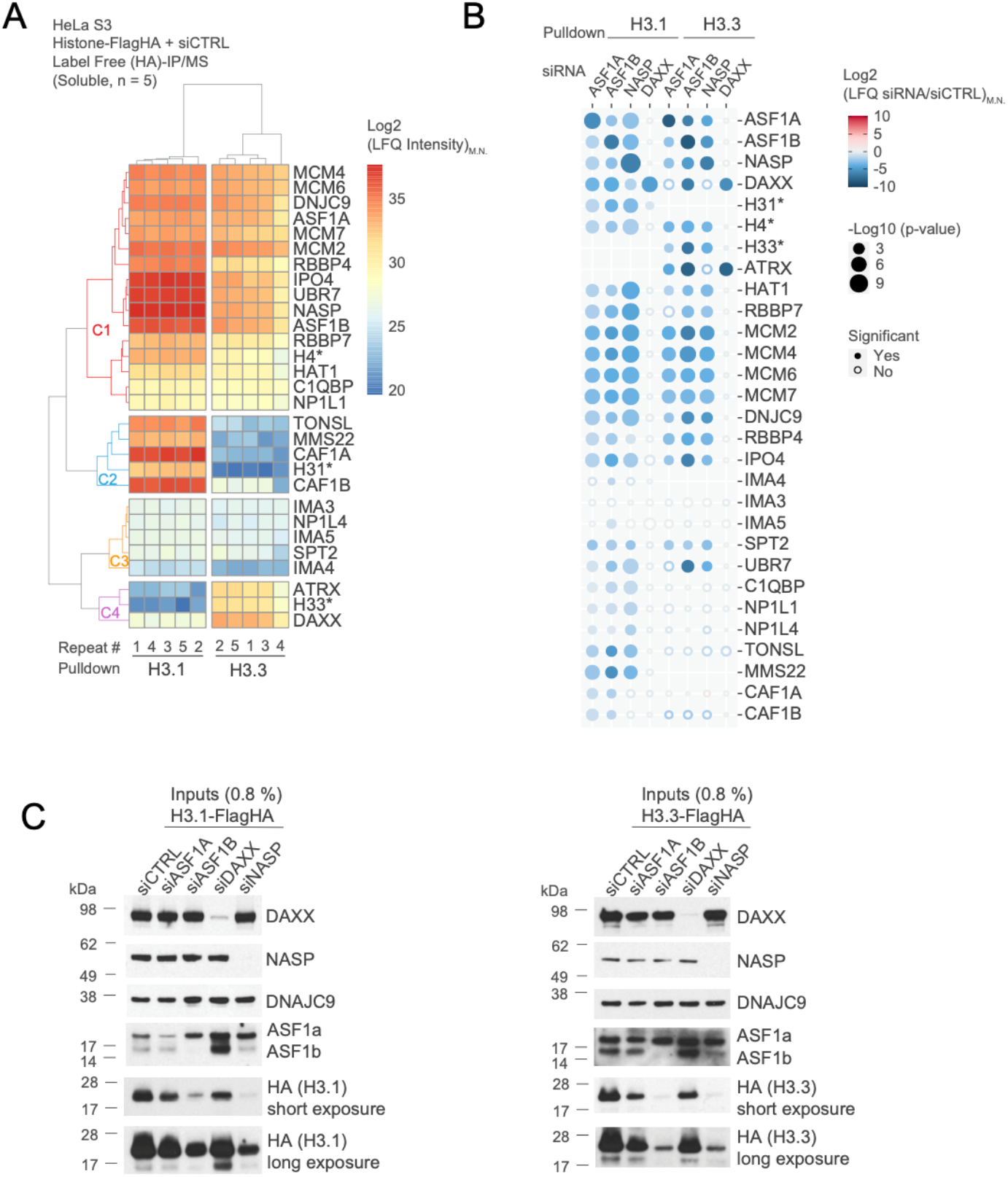
Histone chaperone perturbation demonstrates their functional connectivity. (A) Clustering analysis showing Euclidean distances between median normalized LFQ intensities (LFQ_M.N._) for proteins identified in H3.1 and H3.3 pulldowns by MS. (B) Bubble plot showing the changes in abundance of proteins identified in histone H3.1 and H3.3 pulldowns from extracts siRNA depleted of the chaperones ASF1A, ASF1B, NASP and DAXX compared to control conditions (siCTRL). (A-B) Representative of n=5 biological replicates, with LFQ intensities quantified on a protein or *peptide level by MS. Proteins are referred to by human UniProt protein identification code. See also **Table S1.** (C) Western blot of soluble extract from cells expressing FLAG-HA tagged H3.1 (left) and H3.3 (right) siRNA depleted for ASF1A, ASF1B, DAXX or NASP and compared to control knockdowns siCTRL. Representative of n=5 biological replicates.

First, we performed hierarchical clustering analysis of LFQ intensities for these proteins compared to the peptide-level LFQ intensities for H3.1, H3.3 and H4, to track the abundance of proteins across experiments (**Figure 4A**). As expected H3.1 and H3.3 biological replicates clustered together with histone H3.1 and H3.3 specific peptides validating their respective enrichment. The intensity of histone H4 peptides was more evenly distributed across the H3.1 and H3.3 pulldowns serving as a proxy for factors that bind H3–H4 independently of histone H3 isoform (**Figure 4A**, cluster C1). We observed that the histone chaperones ASF1a/b, DNAJC9, MCM2, RBBP4/7, NASP and NP1L1, along with MCM4/6/7, C1QBP, HAT1, IPO4 and UBR7 clustered together with histone H4, supporting a conserved function for these proteins in both H3.1 and H3.3 supply pathways. Another cluster of proteins, which included IMA3, IMA4, IMA5 and NP1L4 and SPT2, had a similar enrichment profile across conditions. However, their low abundance suggests that they only play a more minor role in soluble H3.1 and H3.3 histone supply (**Figure 4A**, cluster C3). By contrast, DAXX and ATRX formed a distinct cluster with H3.3, interestingly ATRX was stringently selective for H3.3 whereas DAXX was also identified in H3.1 pulldowns albeit to a much lower intensity (**Figure 4A**, cluster C4). The latter suggests an ATRX-independent role of DAXX in H3.1 biology, supported by other observations that DAXX can bind H3.1–H4 (DeNizio et al., 2014; Hammond *et al.*, 2021). Notably, the TONSL-MMS22L complex was only consistently identified in H3.1 pulldowns and clustered with H3.1 peptides similarly to the CAF-1 complex (**Figure 4A**, cluster C2). This argues that TONSL is a H3.1 chaperone in humans, in line with recent findings with the plant homologue TONSUKU (Davarinejad et al., 2022).

We then compared the effects of chaperone knockdowns on these interactions (**Figure 4B**). As previously described (Cook et al., 2011), NASP depletion led to a reduction of the soluble level of both H3.1 and H3.3 (**Figure 4B-C**). Surprisingly, we observed a similar effect with ASF1 depletion, with ASF1b depletion having the strongest effect (**Figure 4B**), despite unperturbed levels of NASP (**Figure 4C**). With the ability of NASP and ASF1 to form a co-chaperone complex, reported here and previously (Campos *et al.*, 2010; Jasencakova *et al.*, 2010), this result suggests that NASP and ASF1 collaborate to protect histones H3–H4 from degradation. Further supporting this idea, the H3 αN helix interaction of NASP required to bridge the co-chaperone complex with ASF1–H3–H4 is also required for H3–H4 stability, whereas loss of the NASP interaction with monomeric H3 is dispensable for this purpose (Bao *et al.*, 2022).

Proteins that bind both H3.1 and H3.3, represented by cluster 1 (**Figure 4A,** cluster C1), were in almost all cases consistently depleted under NASP, ASF1a and ASF1b knockdown conditions (**Figure 4B**). This follows the levels of soluble histones in these conditions, underscoring that ASF1a/b and NASP are master regulators of histone H3.1–H4 and H3.3–H4 supply pathways. Of the lowly abundant factors identified in cluster 3 (**Figure 4A**), only SPT2 followed the same trend (**Figure 4B**), confidently assigning its role in both H3.1 and H3.3 pathways. TONSL, MMS22/MMS22L, C1QBP, NP1L1, and NP1L4 were only lost in pulldowns where H3.1 levels were affected. This was also the case for CAF1A and CAF1B, although surprisingly not in the case of NASP depletion, demonstrating the critical link between ASF1 and the supply of histones H3.1–H4 to CAF1. Meanwhile, DAXX depletion exclusively affected ATRX binding to histone H3.3, without impacting other histone chaperone associations. This argues that DAXX operates as a largely independent arm of the histone chaperone network when depositing histones to heterochromatin and contributes minimally to the supply of histones to other chaperone systems. Otherwise, DAXX and ATRX binding to H3.3 seemed to depend on ASF1 and were most significantly reduced upon loss of ASF1B in H3.3 pulldowns, implying that ASF1b acts partially upstream of DAXX in H3.3 supply to heterochromatin.

### Pre-deposition H3K9 trimethylation of DAXX bound histone H3.3

The concept of a nucleosome assembly pathway dedicated to *de novo* heterochromatin assembly has been proposed based upon the identification of a soluble pool of histone H3 methylated at lysine 9 (H3K9me1 and H3K9me2) (Jasencakova *et al.*, 2010; Loyola et al., 2006; Pinheiro et al., 2012; Rivera et al., 2015). However, it remains unexplored whether and how such pre-modified histones are targeted specifically to heterochromatin. Given the histone dependent association of DAXX with multiple heterochromatin factors, including readers and writers of H3K9me3 (**Figure 2B**), we speculate that DAXX could constitute a likely candidate to mediate targeted silencing by depositing pre-modified histones. To explore this hypothesis, we profiled the PTMs of histone H3–H4 dimers bound to DAXX using MS analysis. For comparison, we included a PTM profile of histone H3–H4 associated with sNASP (**Figure 5A**).

**Figure 5.**
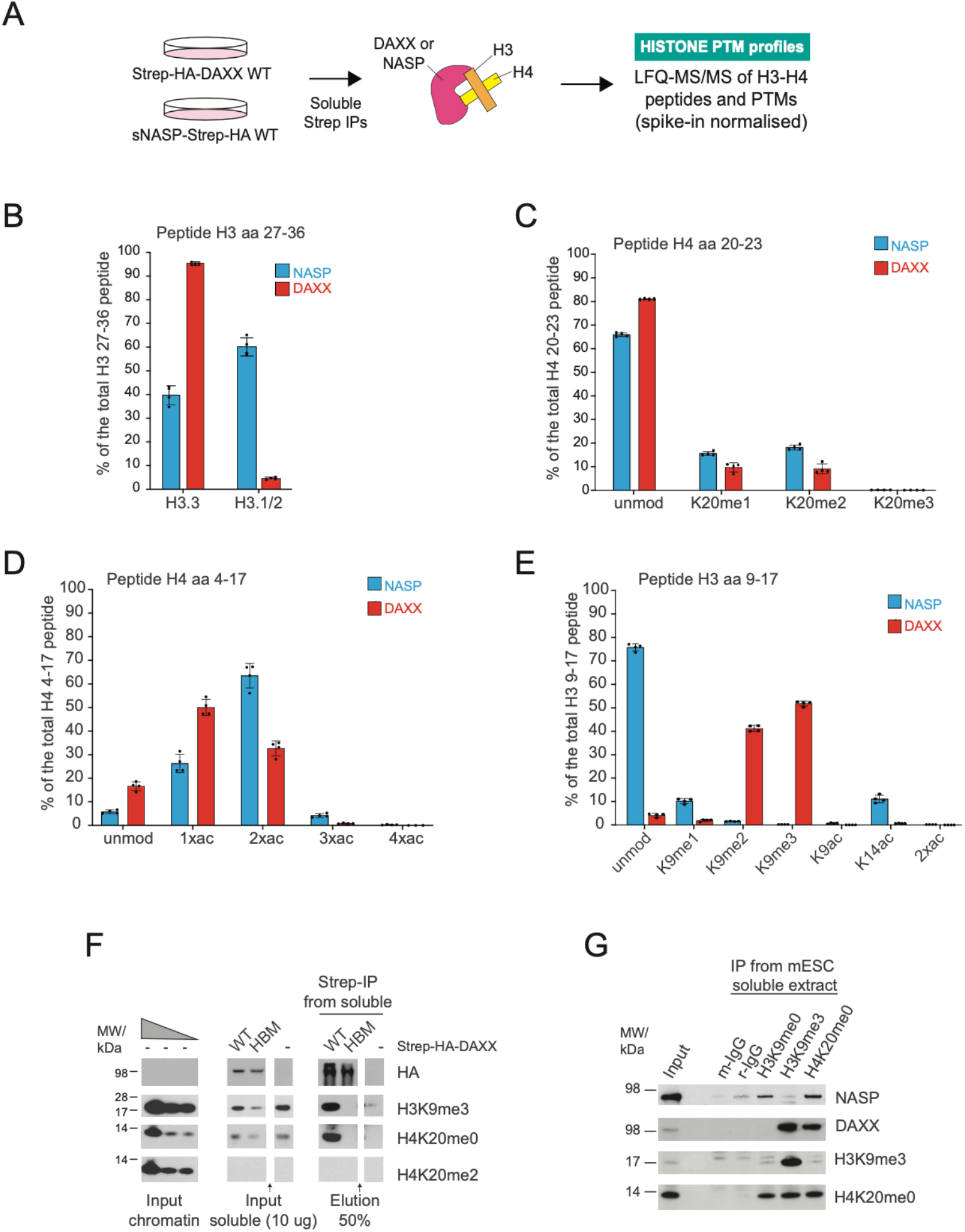
DAXX escorts new H3.3 K9 methylated prior to deposition. (A) Strategy for profiling the peptides and PTMs of histones H3 and H4 from soluble sNASP and DAXX purifications. (B) Quantification of H3.1/2 and H3.3 peptides associated with sNASP and DAXX. (C-E) Quantification of PTMs on H4 peptides 20-23 (top), H4 peptides 4-17 (middle), and H3 peptides 9-17 (bottom) associated with sNASP and DAXX. (B-E) Percentages are relative to the total intensity of related peptides and averaged across n=4 biological replicates of sNASP and DAXX purifications. Error bars represent SD. See also **Figure S4** and **Table S2**. (F) Pull-downs of Strep-HA-tagged DAXX WT or HBM compared with control purifications (−) from soluble cell extracts probed by Western blot for histone modifications and compared to histone PTM levels on chromatin by serial dilution. Representative of n=2 biological replicates. Arrow indicates lanes containing unrelevant samples have been removed. (G) H3K9me0, H3K9me3 and H4K20me0 antibody pulldowns of endogenous histones from soluble mESC extracts compared to an IgG control, probed by Western blot. Representative of n=2 biological replicate.

As expected, DAXX almost exclusively interacted with H3.3 (Drané *et al.*, 2010; Elsässer *et al.*, 2012; Goldberg *et al.*, 2010)(**Figure 5B**), whereas sNASP bound both H3.1/2 and H3.3 (Tagami et al., 2004). Both chaperones were associated with newly synthesized histone H4 (**Figure 5C**), identified by the absence of methylation on H4 lysine 20 (Saredi *et al.*, 2016). Consistently, other PTMs prevalent on nucleosomal histones (Loyola *et al.*, 2006), including H3K4, H3K27, H3K36 and H3K79 methylations (**Figure S4A-C, Table S2**), were not identified on either DAXX or NASP bound histones. Low levels of K14, K18, K23 acetylation were found in DAXX and sNASP complexes, while H3K56ac was not detected (**Figure 5C, Figure S4D-4E**), as expected from previous analysis of ASF1 bound histones (Jasencakova *et al.*, 2010). Di-acetylation of histone H4 on lysine 5 and 14 (H4K5acK12ac) is catalyzed by HAT-1 complex during histone supply (Sobel et al., 1995; Verreault et al., 1998) and serves as another proxy for new histones. We identified di-acetylation of the H4 tail on both DAXX and NASP bound histones (**Figure 5D**), further demonstrating their association with new histones. sNASP associates with the HAT-1 complex during nucleosome assembly (Figure 2A) and ~65 % of sNASP bound H4 were di-acetylated. Since >95 % of histone H4 in complex with ASF1b are di-acetylated (Jasencakova *et al.*, 2010), our data supports that sNASP and the HAT-1 complex are upstream of ASF1 in the histone supply chain (Campos *et al.*, 2010). H4 di-acetylation was even lower in the DAXX complex (~30 %, **Figure 5D**), which is perhaps the result of DAXX associating with the histone deacetylase complex members SAP18 (**Figure 1B**) and HDAC1/2 (Hoelper *et al.*, 2017; Hollenbach et al., 2002).

Strikingly, ~ 90% of histone H3 in complex with DAXX was di- or tri-methylated on H3K9, while sNASP mainly associated with histones unmethylated at H3K9 (**Figure 5E**), similar to ASF1b (Jasencakova *et al.*, 2010). We consolidated the finding that DAXX associates with newly synthesized histones marked with both H4K20me0 and H3K9me3 pre-deposition by western blotting (**Figure 5F**). H3K9me3 has previously been detected in endogenous DAXX complexes purified from mouse embryonic stem cells (mESC) (Elsässer *et al.*, 2015), we confirmed this and further established that endogenous DAXX binds new histones demarked by H4K20me0 (**Figure 5G**). Collectively, this demonstrates that DAXX binds newly synthesized non-nucleosomal histones H3.3–H4 carrying H3K9me3 prior to their deposition on DNA (**Figure 5B-G)**, identifying a conserved DAXX-centered pathway for *de novo* heterochromatin assembly.

### DAXX stimulates H3K9me3 methyltransferase activity

To decipher the mechanism of H3K9 trimethylation in DAXX complexes, we assessed the contribution of key H3K9 methyltransferases SETDB1 and SUV39H1/2, previously shown to associate with DAXX (He *et al.*, 2015; Hoelper *et al.*, 2017), on DAXX bound H3K9 methylation levels. Depletion of either SETDB1 or SUV39H1/2 caused a significant reduction in the H3.3 K9me3 levels associated with DAXX (**Figure 6A, Figure S5A, Table S2**), with SETDB1 depletion having the strongest effect. This data suggests that SETDB1 may more efficiently covert H3K9me2 to K9me3 compared to SUV39H1/2, as previously suggested (Loyola *et al.*, 2006). The fact that DAXX carries both H3.3 K9me2 and K9me3 (**Figure 5C**) and interacts with readers and writers of K9 methylation (**Figure 1A and 2A**), implicates DAXX in handling histones during the catalysis of H3.3 K9me3. In support of this hypothesis, we found that DAXX potentiates the activity of SETDB1 towards K9 trimethylation of H3.3–H4 dimers *in vitro* (**Figure 6B**). This ability was not paralleled by ASF1b (**Figure 6B**), supporting a unique ability of DAXX to stimulate the catalysis of H3.3 K9me3 on H3–H4 dimers.

**Figure 6.**
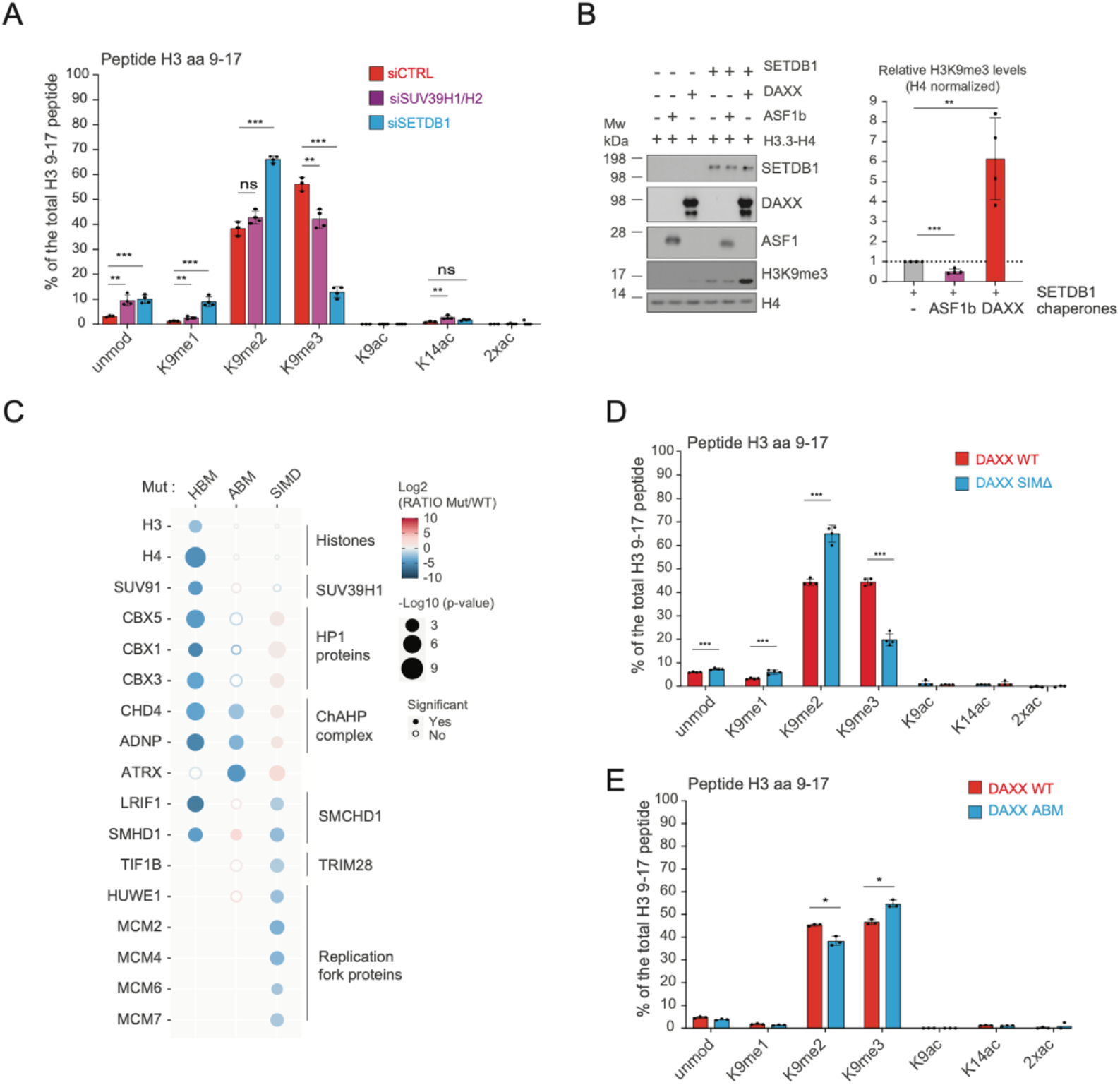
DAXX promotes the catalysis of H3.3 K9me3 in collaboration with SETDB1. (A) Quantification of PTMs on H3 peptides 9-17 associated with DAXX upon SETDB1, SUV39H1/H2 or control siRNA depletions averaged across n=4 (siSETDB1 and siSUV39H1/H2) and n=3 (siCTRL) biological replicates. P values represent unpaired two-sided t-tests (from left to right unmod: 0.004146, 0.000955; K9me1: 0.007882, 0.000913; K9me2: 0.081576, 0.000012; K9me3: 0.002794, 0.000003; K9ac: 0.185665, 0.005248; K14 ac: 0.010470, 0.009786). See also **Figure S5A**. (B) In vitro histone methyltransferase assay with recombinant proteins analysed by Western blot with average quantification of H3K9me3 representative of n=4 independent experiments. Error bars represent SD. P values represent unpaired two-sided t-tests (from left to right H3K9me3 quantification: P=0.0002; 0,0024) (C) Bubble plot showing the changes in abundance for factors associated with DAXX HBM, ABM or SIMΔ mutants compared to WT DAXX. Proteins are referred to by human UniProt protein identification code. Data generated from n=4 biological replicates. See also **Table S1** and **Figure S5D-E**. (D) Quantification of PTMs on H3 peptides 9-17 associated with DAXX WT or SIMΔ mutant averaged across n=4 biological replicates. P values represent unpaired two-sided t-tests (from left to right unmodified=0.000707; K9me1=0.000982; K9me2=:0.000034; K9me3=0.000003; K9ac=0.316984; K14 ac=0.458609). (E) Quantification of PTMs on H3 peptides 9-17 associated with DAXX WT or ABM mutant averaged across n=3 biological replicates. P values represent unpaired two-sided t-tests (K9me2=0.003661; K9me3=0.002382) (A and D-E) Percentages are relative to the total intensity of related peptides and averaged across biological replicates of DAXX purifications. Error bars represent SD. See also **Table S2**.

Whilst SUV39H1/2 is a histone dependent interactor of DAXX, we were unable to identify SETDB1 in DAXX interactomes (**Figure 1–2**), contrasting previous observations (Hoelper *et al.*, 2017). Considering that the association of DAXX with ATRX and SUMOylated proteins can drive DAXX-mediated transcriptional silencing (Dyer *et al.*, 2017; Hoelper *et al.*, 2017; Lin et al., 2006), we speculated that SETDB1 recruitment could be transient and mediated by ATRX or SUMOylation. Therefore, we profiled the dependency of the DAXX bound factors on ATRX and SUMOylation and assayed the levels of histone PTMs associated with each mutant. Based on previous studies (Escobar-Cabrera et al., 2011; Hoelper *et al.*, 2017), we generated ATRX binding mutant (ABM) and deleted two annotated SUMO interacting motifs (SIMs) to generate a SIM mutant (SIMΔ) (**Figure S5B-C**).

Comparison of the interactomes of DAXX ABM and SIMΔ mutants to the WT and HBM showed an intricate dependency network of factors recruited by DAXX (**Figure 6C, Figure S5D-E**). For example, DAXX interaction with members of the ChAHP chromatin remodeling complex (ADNP, CHD4) (Ostapcuk et al., 2018) was lost for DAXX ABM (**Figure 6C, Figure S6C**). Similarly, DAXX association with the heterochromatin proteins 1 (HP1) was reduced in the ATRX binding mutant (**Figure 6C, Figure S6C**). This corroborates ATRX binding to HP1 proteins (Lechner et al., 2005) and ADNP (Teng *et al.*, 2021) and suggests ATRX mediates DAXX-H3.3-H4 recruitment to distinct nuclear complexes. Deletion of the SIM domains in DAXX reduced the stability of the mutant compared to the WT protein (**Figure S5E**), showing that SUMO binding is important for the stability of DAXX. However, the loss of SUMO binding did not affect the association of DAXX with H3 and H4, indicating that DAXX SIMΔ retained the ability to chaperone histones (**Figure 6C, Figure S5E)**. Upon bait normalization, our results revealed that DAXX forms a substantial number of SUMO dependent interactions (**Figure S5E**), which reflect the SUMO dependent recruitment of DAXX to subnuclear compartments including PML bodies (Corpet *et al.*, 2020; Lin *et al.*, 2006). We identified SUMOylation as a mechanism for DAXX to connect with SMCHD1/SMHD1, LRIF1, TIF1B/TRIM28/KAP1 and replication fork proteins (MCM2-7, HUWE1) (**Figure 6C**), which are identified as SUMOylated proteins (Hendriks et al., 2014) and known or putative SETDB1-linked factors (Ichihara et al., 2021; Keniry et al., 2016; Nozawa et al., 2013; Schultz et al., 2002; Teng *et al.*, 2021). These factors were either marginally enriched with DAXX ABM (TIF1B, SMCHD1, LRIF1 and HUWE1) or not affected (MCM2-7). In contrast, the ChAHP complex and HP1 proteins were trapped by the SIMΔ mutant (**Figure 6C, Figure S5D-E**).

Strikingly, the DAXX SIMΔ mutant reduced the DAXX associated H3K9me3 levels (**Figure 6D, Figure S5E, Table S2**) to a similar extent as the loss observed upon depletion of SETDB1 (**Figure 6A**). This supports that the SUMO-dependent recruitment of SETDB1-linked factors to DAXX is associated with the catalytic mechanism of H3K9me3 methylation (**Figure 6C**). Consistent with this, treatment with the SUMO activating-enzyme inhibitor ML-792 (He et al., 2017) also reduced H3K9 trimethylation on DAXX bound histones (**Figure S5G**). Meanwhile, loss of ATRX caused a slight increase in the levels of H3K9me3 with DAXX (**Figure 6E, Table S2**), consistent with the slightly higher level of SUV39H1 and TRIM28 associated with this mutant (**Figure 6C**). This demonstrates that ATRX is not involved in the DAXX-dependent establishment of H3K9me3 on soluble histones. Meanwhile, both ATRX and SUMO binding coordinate DAXX interactions with discrete heterochromatin complexes (**Figure 6C**), potentially orchestrating *de novo* H3.3 K9me3 deposition at alternative chromatin sites (**Figure 7**).

**Figure 7.**
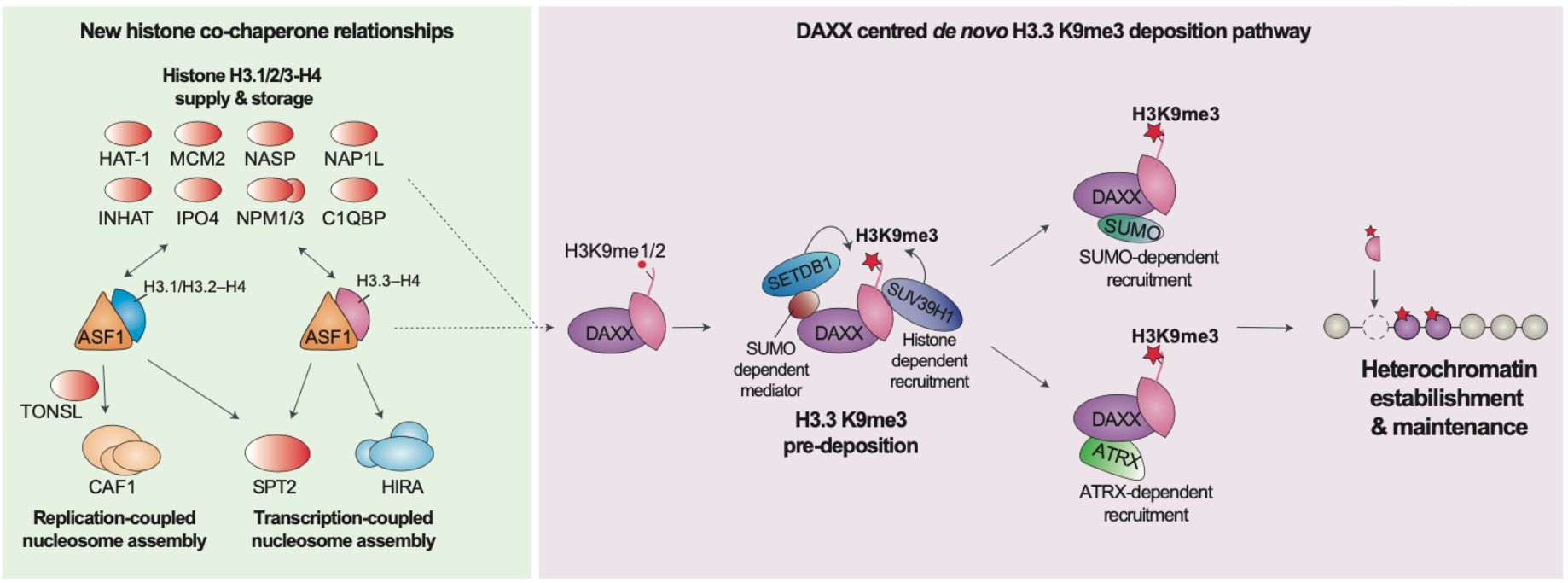
Model. During histone supply ASF1 handles H3.1/2/3–H4 dimers and forms several histone dependent co-chaperone complexes with other histone chaperones (colored in red), notably ASF1 channels H3 variants towards distinct deposition complexes on chromatin. We identified a new ASF1-centred histone supply pathway to SPT2 that is H3 variant independent, as well as an upstream role for ASF1 in delivering H3.3 histones to DAXX, and a H3.1 variant specificity in TONSL. We found DAXX facilitates the catalysis of H3.3K9me3 through SETDB1 and SUV39H1 methyltransferase recruitment pre-deposition. SETDB1-mediated H3K9me3 is driven by SUMO-dependent interactions with DAXX (e.g., TRIM28 or PML), and we speculate DAXX can be recruited via ATRX or SUMO for H3K9me3 histone deposition in different genomic contexts supporting *de novo* heterochromatin silencing.

## Discussion

In this work, we provide a comprehensive interactome analysis of the histone chaperones central to H3 variant supply pathways. Through this, we discover unexplored histone dependent connections with other cellular processes and reveal unique functionalities contributed by each chaperone to the global histone chaperone network. In addition, we identify a mechanism for targeted *de novo* heterochromatin formation through dissection of the DAXX assembly pathway. This valuable resource strengthens our understanding of histone chaperone biology as a whole and serves to highlight how these remarkable proteins collaborate within a cellular context.

We expand the central role of ASF1 in histone supply pathways, identifying uncharacterized histone dependent co-chaperone complexes with SPT2, C1QBP, NPM1-3 and NAP1L1/4 proteins (**Figure 7**). SPT2 and NAP1 bind (H3-H4)_2_ tetramers (Bowman et al., 2011; Chen *et al.*, 2015), while the binding mode of C1QBP and NPM1-3 remains to be established. Predicting the molecular basis for SPT2 co-chaperoning a H3–H4 with ASF1, we find that the co-chaperone complex contains H3–H4 dimers. This is reminiscent of MCM2 and Vps75, which can both bind (H3-H4)_2_ tetramers alone and form co-chaperone complexes with ASF1 and dimeric H3–H4 (Bowman *et al.*, 2011; Hammond *et al.*, 2016; Huang *et al.*, 2015). SPT2 and HIRA are both linked to transcription (Chen *et al.*, 2015; Goldberg *et al.*, 2010; Osakabe *et al.*, 2013), but whilst HIRA is H3.3 specific (Tagami *et al.*, 2004), our data reveal that SPT2 binds both H3.1 and H3.3. This supports a collaboration between ASF1 and SPT2 in handling both H3.1 and H3.3 during transcription (**Figure 7**). Given that H3.1 is generally not deposited *de novo* in a transcription-coupled manner, we envision that ASF1 participates with SPT2 in handling eviction and recycling of both the canonical and replacement H3 variants, possibly allowing them to re-enter the supply chain. In this respect, ASF1 has been implicated in the recycling of histone during transcription in yeast (Schwabish and Struhl, 2006) and mammals (Torné et al., 2020), and ASF1–SPT2 cooperation could be required during this process.

Our interrogation of the histone chaperone network also revealed a H3.1 specificity of TONSL-MMS22L, which has important implications for the mechanism of marking post-replicated chromatin for error-free DNA repair. In this respect, TONSL would read H4 K20me0 to specify new histones (Saredi *et al.*, 2016) and H3.1 to specify delivery to chromatin via a DNA-replication-coupled mechanism (**Figure 7**). This may be required to restrict TONSL-MMS22L function in homologous recombination to post-replicative chromatin, as replication-independent deposition of new H3.3–H4 carrying H4K20me0 would not support TONSL-MMS22L recruitment. Notably, the plant homologue TONSUKU which is also specific for H3.1 (Davarinejad et al., 2022) lacks the ankyrin repeat domain required for H4 K20me0 recognition (Davarinejad et al., 2022; Saredi *et al.*, 2016). Together, this suggests TONSL-MMS22L has post-replicative functions not mirrored by the plant counterpart.

We identified DAXX as an independent arm of the histone chaperone network that stimulates catalysis of H3K9me3 on H3.3–H4 dimers prior to their deposition onto DNA. This provides a mechanism for assembly of heterochromatin *de novo* through the targeting of DAXX to different regions of the genome, and helps to explain why DAXX-mediated H3.3 deposition is required for H3K9me3-mediated silencing of repetitive DNA elements and imprinted regions (Elsässer *et al.*, 2015; He *et al.*, 2015; Voon *et al.*, 2015). We show that SETDB1 and SUV39H1 methylate DAXX bound H3.3-H4 dimers, however whilst the depletion of SETDB1 and SUV39H1/2 reduces H3K9me3 levels bound by DAXX the H3K9me1/2 levels were not decreased. This suggests that other methyltransferases act redundantly with or upstream of SETDB1 and SUV39H1/2. SETDB1 is also responsible for the catalysis of K9me1/2 on a fraction of new histone H3 during translation (Rivera *et al.*, 2015) and H3K9me1/2 marking of new histones has previously been proposed to potentiate heterochromatin assembly (Loyola *et al.*, 2006; Loyola *et al.*, 2009; Pinheiro *et al.*, 2012). While the mechanism underlying targeted deliver of these pre-marked histones remains unclear, we envision they could represent H3.3 destined for the DAXX assembly pathway. In addition, ~5% of ASF1b bound histones carry H3K9me1 in the S phase (Jasencakova *et al.*, 2010), which could represent the fraction of histones delivered to DAXX for conversion to H3K9me3 and deposition. In this respect we found the supply of histones H3.3-H4 to DAXX to be dependent on ASF1b (**Figure 7**), and a similar collaborative link has been identified between the DAXX-Like Protein (DLP) and ASF1 in flies (Fromental-Ramain et al., 2017).

While SUV39H1 interacts with DAXX in a histone dependent manner, we propose that SETDB1 recruitment to DAXX is mediated by SUMOylation via bridging factors, such as TRIM28 and PML bodies (**Figure 7**). In this respect, auto-SUMOylation of TRIM28 was found to be important for the association with SETDB1 (Ivanov et al., 2007; Zeng et al., 2008) and both DAXX and SETDB1 localizes at the PML bodies in a SUMO dependent manner (Cho et al., 2011; Corpet *et al.*, 2020; Ishov et al., 1999; Lin *et al.*, 2006). Future structural investigations will be required to address how DAXX positions the H3 tail for methylation by SETDB1/SUV39H1, however the significant reorientation of the H3 αN helix observed in the DAXX complexes (Elsässer *et al.*, 2012) could be at the heart of why DAXX stimulates H3.3K9me3 catalysis and ASF1 does not. The DAXX-bound H3.3–H4 supply pathway diverge into ATRX- and SUMO-dependent delivery pathways (this work and (Hoelper *et al.*, 2017)), likely reflecting H3K9me3 deposition at distinct chromatin locations (**Figure 7**). For instance, recruitment of ATRX– DAXX–H3.3K9me3–H4 to the ChAHP chromatin remodeling complex could target ADNP binding sites for H3K9me3-mediated silencing. Consistent with this model, ADNP binding sites in euchromatin show moderate enrichment for H3K9me3 (Ostapcuk et al., 2018). We propose that the DAXX-centred supply of H3.3–H4 marked with H3K9me3 provides a means to target heterochromatin formation to diverse sites (**Figure 7**), contributing to *de novo* H3K9me3 establishment at stalled replication forks (Teng *et al.*, 2021), silencing of viral genomes after infection (Cliffe and Knipe, 2008; Cohen et al., 2018; Tsai et al., 2014), and H3K9me3 maintenance at repetitive elements counter-acting the dilution of H3K9me3-marked chromatin during DNA replication.

Collectively, our work presents a panoramic view of the histone chaperone network and elucidates the molecular basis for DAXX-mediated heterochromatin assembly. These discoveries demonstrate the pivotal importance of histone chaperone-centered histone supply pathways in the support of chromatin functionality, which has broad implications for the epigenetic regulation of biological systems.

### LIMITATION OF THE STUDY

Our proteomics and network analysis integrated the interactome of seven different H3–H4 histone chaperones. However, additional H3–H4 histones chaperones (e.g. FACT, SPT2, IPO4) were not directly targeted in our analysis. Our work should therefore serve as a crucial framework for understanding the cooperation between histone delivery systems that can be further expanded in the future. Other approaches, such as proximity labelling proteomics (Bio-ID or APEX2) may also help to capture transient interactions we may have missed in our experimental strategy. There is also the potential for the non-histone dependent and histone-dependent interactors that we have identified to be mediated by RNA or other proteins, therefore reconstitution-based and structural approaches will help to resolve details of the interactomes we have provided.

## ACKNOWLEDGMENTS

We thank Nadezhda T. Doncheva and Lars Juhl Jensen for help with Cytoscape; A.G. and M.L.N. lab members for helpful discussions. M.L.N. is supported by the Independent Research Fund Denmark (0135-00096B and 8020-00220B), the European Union’s Horizon 2020 research and innovation program under grant agreement EPIC-XS-823839, and the Danish Cancer Society (R146-A9159-16-S2). A.G. is supported by the European Research Council (ERC CoG 724436), the Lundbeck Foundation (R198-2015-269 and R313-2019-448), and the Independent ResearchFund Denmark (7016-00042B). Research at CPR is supported by the Novo Nordisk Foundation (NNF14CC0001).

## AUTHOR CONTRIBUTIONS

A.G., M.C. and C.M.H. conceived the study, designed experiments and wrote the manuscript with input from all authors. M.C. led the interactome characterization of the histone chaperones and DAXX-mediated histone supply, including generating cell lines, plasmids, pull-downs, Western Blot, SILAC and label-free MS, histone PTM analysis and in vitro methylation assay. M.C. performed the final proteomics data analysis in consultation with I.A.H. C.M.H. performed label-free histone pull-downs and related data analysis. AlphaFold predictions were performed by C.M.H. in consultation with M.W. and G.M. I.A.H. and J.D.E. optimized and performed MS sample cleanup, designed MS methodology, acquired the majority of SILAC and label-free MS data, and performed preliminary MS data analysis, all in consultation with M.L.N. M.V.A. and V.S. acquired and analyzed histone PTMs MS in consultation with A.I. C.S. acquired mass spectrometry data in consultation with J.R.

## DECLARATION OF INTERESTS

C.M.H. and A.G. are inventors on a patent covering the therapeutic targeting of TONSL for cancer therapy. A.G. is cofounder and chief scientific officer (CSO) of Ankrin Therapeutics. A.G. is a member of Molecular Cell’s scientific advisory board. A.I. and M.V.A. are cofounders of EpiQMAx.

## STAR Methods

### RESOURCE AVAILABILITY

#### Lead Contact

Further information and requests for resources and reagents should be directed to and will be fulfilled by the Lead Contact, Anja Groth.

#### Materials Availability

All stable and unique reagents generated in this study are available from the Lead Contact subject to a Materials Transfer Agreement.

#### Data and Code Availability

Histone PTMs mass spectrometry datasets that support the findings have been deposited to the ProteomeXchange Consortium via the PRIDE (Perez-Riverol et al., 2019) with the accession code: PXD034924.

The Immunoprecipitation-mass spectrometry proteomics data have been deposited to the ProteomeXchange Consortium via the PRIDE partner repository with the dataset identifier PXD034888.

### Experimental model and subject details

#### Cell lines

##### Cell line generation and transfection

H3.1 (HeLa S3 pLVX-TetOne-puro H3.1-FlagHA), H3.3 (HeLa S3 pLVX-TetOne-puro H3.3-FlagHA) and control (HeLa S3 pLVX-TetOne-puro-TwinStrep-HA) cell lines were published previously – see Key Resource Table. Cell lines expressing ASF1a, ASF1b, sNASP, HJURP and DAXX (WT and mutants) from pLVX-TetOne-puro constructs were created by lentiviral transductions of HeLa S3 suspension cells followed by 24 hours of puromycin selection (1 μg/ml). The lentivirus-containing media were collected and filtered using a 0.45 μm syringe 60 hours after transfecting of 293FT cells with 5 μg pVSV, 8 ug psPAX2 and 10 μg pLVX-TetOne plasmids using Lipofectamine 2000 reagents according to the manufacturer’s instructions. The presence of lentiviral particles was confirmed using Lenti-X GoStix Plus according to the manufacturer’s instructions. All used cell lines generated in this study tested negative for mycoplasma contamination. HeLa S3 and 293FT cell lines were derived from female subjects.

##### Cell culture

HeLa S3 and 293FT cells were grown in DMEM + GlutaMAX (Thermo Fisher Scientific) medium supplemented with 10% FBS (Hyclone) and 1 % penicillin and streptomycin. E14 mESCs (male) were grown in on plates coated with 0.2% gelatin (Sigma, G9391) in DMEM media (GIBCO, 10829018) supplemented with GlutaMAX-pyruvate (Thermo Fisher Scientific) with fetal bovine serum (15 %, Hyclone), LIF (made in-house), 1x non-essential amino acids (Gibco), 1x penicillin/streptomycin (Gibco) and 2-beta-mercaptoethanol (0.1 μM). E14 mESCs were passaged using Trypsin-EDTA (Gibco). All cells were grown in a humidified incubator at 37 °C with 5% CO_2_. HeLa S3 cell lines expressing pLVX-TetOne-Puro cell lines were grown under Puromycin selection (1 µg/ml) and the expression of the chaperones (ASF1a, ASF1b, sNASP, HJURP or DAXX) and histones (H3.1, H3.3) was induced by treatment with 2 µg/ml Doxycycline for 24-36 hours. For SILAC experiments cells were grown in RPMI 1640 Medium for SILAC (Thermo scientific, 88365) supplemented with dialyzed FBS (Thermo Scientific), MEM non-essential amino acid mix (Thermo Scientific), GlutaMax (Thermo Scientific), and isotopically labelled arginine (316 μM) and lysine (547 μM). triple SILAC experimental conditions employed heavy Lys8-Arg10, medium Lys4-Arg6, or light Lys0-Arg0. The HeLa S3 pLVX-TetOne-puro-TwinStrep-HA control cell line for all triple SILAC experiments was cultured with light amino acids Arg0 and Lys0 (A6969 and L8662, Sigma) in all biological replicates, while the HeLa S3 expressing WT and mutant chaperones were label-swapped between medium Arg6 and Lys4 (CNLM-2265-H1 and DLM-2640-1, Cambridge Isotope Laboratories) and heavy Arg10 and Lys8 (CNLM-539-H1 and CNLM-291-H-1, Cambridge Isotope Laboratories) amino acid pairs across biological replicates. Where indicated, cells were treated with 1 µM SUMO1/2 inhibitor ML-792(He et al., 2017) for 6 hours. For siRNAs depletion experiments, cells were transfected with Lipofectamine RNAiMAX (Invitrogen) and 20 nM of Silencer^®^ Select siRNA for 96 hours, prior to being harvested as per the manufacturer’s instructions.

### Method details

#### Plasmid generation

The TwinStrep-HA tag (sequence TGGGGSGGGASWSHPQFEKGGSGGGSWSH PQFEKGGYPYDVPDYA*) was synthesised and cloned (by Genscript) between the EcoRI and BamHI sites of the pLVX-TetOne-Puro vector (631849, Clontech). Subsequently, ASF1a, ASF1b, sNASP, HJURP and DAXX coding sequences were sub-cloned into pLVX-TetOne-puro-TwinStrep-HA by amplifying their respective chaperones cDNA (from Origene plasmids) with primers that also create the homologous arms and using these PCR products as “mega-primer” pairs. All the chaperone cDNAs were cloned in frame with an N-terminal of TwinStrep-HA tag, except for sNASP which was TwinStrep-HA tagged at the C-terminal. The chaperones mutations (HBMs, ABM) were performed using site directed mutagenesis of the respective pLVX-TetOne-puro-Chaperone-WT-TwinStrep-HA plasmids. Site directed mutagenesis was performed using established QuickChange mutagenesis protocols (Stratagene) or Infusion HD-directed mutagenesis (Takara). For Infusion HD-directed mutagenesis template plasmids were amplified with Phusion HF (F530S, Thermo Scientific) using mutagenic primers that also created homologous arms which, after PCR purification (28104, QIAgen) and Dpn1 digest (R0176L, NEB), were recombined through Infusion HD cloning (638933, Takara). The DAXX mutations for the and SIMΔ plasmid was performed by Genescript.

#### Cell extracts

Soluble extracts were prepared by washing the cells twice with cold PBS and pelleting them by centrifugation (300 g, 3 mins) at 4 °C. The cell pellet was resuspended in ice cold NP40-NaCl buffer (300 mM NaCl, 0.05 % Nonidet P40, 50 mM Tris-HCl pH 7.6, 0.1 mM EDTA, 5 % glycerol) with freshly added inhibitors (NaF (5 mM) and β-Glycerolphosphate (10 mM), Phenylmethanesulfonyl fluoride (0.1 mM), Leupeptin (10 µg/ml), Pepstatin A (10 µg/ml), Trichostatin A (100 ng/ml), Na3VO4 (0.2 mM)) and left 15 minutes at 4 °C. Subsequently, the lysate was cleared by centrifugation (11,000 g, 20 min), transferred to a new tube, centrifuged again (11,000 g, 10 min) and filtered (0.45 μm). For extracting chromatin bound proteins, the pellets derived from the soluble extraction were digested for 1 hour at 37 °C with 0.015 volumes of 25 U/μl Benzonase (Millipore, 70746) in 1 volume NP40-NaCl buffer supplemented with 0.01 volumes of 1 M MgCl2. The resultant chromatin extracts were cleared by centrifugation (16,000 g, 3 min, 4 °C), and supernatants were transferred to new tubes. The resultant soluble and chromatin fractions were used directly for immunoprecipitation experiments, western blot analysis, or otherwise stored at −80 °C.

#### Immunoprecipitation and Western blot analysis

Protein concentrations were measured using Pierce™ 660 nm Protein Assay Reagent (Thermo Scientific) and equalized using NP40-NaCl extraction buffer. For immunoprecipitation of tagged proteins, Strep-HA-chaperone and histone-Flag-HA extracts were incubated with MagStrep “type3” XT beads (2-4090-010, iba) or anti-HA (26181, Thermo scientific), respectively, for 3 hours at 4 °C. After incubation, chaperone-IP beads were washed twice using ice-cold wash buffer (150 mM NaCl, 0.02% Nonidet P40, 50 mM Tris-HCl pH 7.6, 0.1 mM EDTA, 5% glycerol), and additionally washed four times with ice-cold wash buffer lacking glycerol and NP-40, prior to elution with 1X Strep-Tactin^®^XT elution buffer (BXT buffer, 2-1042-025, Iba) at RT for 1 h. SILAC-labeled samples were subjected to in-solution tryptic digestion, while samples were probed by western blot analysis. Histone IPs from siRNA treated cell extracts were washed exclusively in NP40-NaCl buffer and then additionally washed with minimal wash buffer (MWB: 300 mM NaCl, 50 mM Tris pH 7.6) and NH4HCO3 (50 mM) prior to on-bead tryptic digestion, following the same protocol as previously described (Hammond *et al.*, 2021). For immunoprecipitation of endogenous H3K9me0 and H3K9me3, 4 mg of soluble extracts were pre-cleared using 50 µL of BSA-blocked (10 mg/l BSA in 1.5 mM TRIS, pH 7.6) Protein A-agarose beads (20333, Thermo Scientific) for 40 minutes at 4 °C, followed by incubation with 50 μL of antibody-coupled BSA-blocked beads (2 μg of H3K9me3 or H3K9me0 antibodies) for 4 hours at 4 °C. Control rabbit IgG (2 μg, Cell Signalling Technology, 2729) and mouse IgG2a (2 μg, ab18415, Abcam) were coupled for the control pull-downs. The beads were then washed five times with ice-cold buffer (150 mM NaCl, 0.02% Nonidet P40, 50 mM Tris-HCl pH 7.6, 0.1 mM EDTA, 5% glycerol), and protein complexes were eluted using Laemmli sample buffer (50 mM Tris-HCl pH 6.8, 1% SDS, 10% glycerol, 25 mM DTT) for 20 minutes at 98 °C. Western blots from four independent biological replicates were quantified using the software ImageJ, version 1.0.

#### In vitro histone methyltransferases assay

100 nM of (H3.3–H4)_2_ tetramer (Reaction biology, HTM-14-438) were incubated with equimolar concentration of either recombinant DAXX (Origene, TP326603) or ASF1b (Abcam, ab130033) in a reaction system containing 50 mM TRIS-HCL pH 8, 0.02% Triton X-100, 2 mM MgCl2, 1 mM TCEP, 50 mM NaCl, and 10% glycerol, for 1 hour at RT. The reaction system was then supplemented with 2 nM of recombinant SEDTB1 (Active Motif, 3152) and 50 μM of SAM (Bionordika, NEB-B9003S) for 15 minutes at RT. H3K9me3 (dilution 1:1000, Abcam, ab176916) antibody was used to detect the reaction product in a western blot assay. H3K9me3 bands were quantified and normalized to H4 and shown relative to SETDB1 activity without chaperones in ImageJ.

#### Antibodies

Western blots were performed with the following antibodies: DNAJC9 (1:1000, ab150394, Abcam), HA (1:3000-5000, C29F4 #3724, Cell Signaling Technology), DAXX (1:250, HPA008736, Sigma), H3 (1:500, ab10799, Abcam), H4 (ab17036, Abcam), Tubulin (1:10000, ab6160, Abcam), H3K9me3 (1:1000, ab194296, Abcam), H3K9me0 (1:1000, 91155, Active Motif), NASP (1:1000, ab181169, Abcam), H4K20me0 (1:500, ab227804, Abcam), H4K20me2 (1:500, C15200205, Diagenode), Actin (1:5000, A5316, Sigma), SUV39H1 (1:500, 8729, Cell Signaling Technology), SUV39H2 (1:500, ab107225, Abcam), SETDB1 (1:1000, ab107225, Abcam), TRIM28 (1:500, 4124, Cell Signaling Technology), ASF1 (1:1000), ADNP (1:500, Bethyl), SPT2 (1:1000, Abcam), UBR7 (1:1000, Bethyl), CBX3 (1:500, Abcam), ERCC6/ERPG3 (1:200 Santa Crutz).

#### MS sample preparation

Samples were digested using sequencing-grade modified trypsin, either in-gel, in-solution, or on-beads, according to standard procedures. In case of on-bead digestion, peptides were 0.45 µm filtered, and cysteine residues were reduced and alkylated by concomitantly adding tris(2-carboxyethyl)phosphine (TCEP) and chloroacetamide (CAA) to a final concentration of 5 mM for 30 min at 30 °C. Preparation of StageTips (Rappsilber et al., 2003), and high-pH cleanup of samples on StageTip, was performed essentially as described previously (Hendriks et al., 2018). Quad-layer StageTips were prepared using four punch-outs of C18 material (Sigma-Aldrich, Empore™ SPE Disks, C18, 47 mm). StageTips were equilibrated using 100 μL of methanol, 100 μL of 80% ACN in 200 mM ammonium hydroxide, and two times 75 μL 50 mM ammonium. Samples were supplemented with 1/10^th^ volume of 200 mM ammonium hydroxide (pH >10), just prior to loading them on StageTip. The StageTips were subsequently washed twice with 150 μL 50 mM ammonium hydroxide, and afterwards eluted using 80 μL of 25% ACN in 50 mM ammonium hydroxide. All fractions were dried to completion in protein-LoBind tubes (Eppendorf), using a SpeedVac for 2 h at 60°C, after which the dried peptides were dissolved using 11 μL of 0.1% formic acid, and stored at −20 °C until MS analysis.

#### Sample preparation for histone modification analysis by MS

Sample preparation and MS analysis were performed according to the EpiQMAx GmbH protocols. Briefly, protein eluted from histone chaperones pulldowns were resuspended in Lämmli buffer and separated by a 14-20% gradient SDS-PAGE, stained with Coomassie (Brilliant blue G-250). Protein bands in the molecular weight range of histones (15-23 kDa) were excised as single band/fraction. Gel slices were destained in 50% acetonitrile/50mM ammonium bicarbonate. SIL heavy standards were spiked in at a concentration of 166 fmoles each. Lysine residues were chemically modified by propionylation for 30 min at RT with 2.5% propionic anhydride (Sigma) in ammonium bicarbonate, pH 7.5. Subsequently, proteins were digested with 200ng of trypsin (Promega) in 50mM ammonium bicarbonate overnight and the supernatant was desalted by C18-Stagetips (reversed-phase resin) and carbon Top-Tips (Glygen) according to the manufacturer’s instructions. After desalting, the eluent was speed vacuumed until dryness and stored at −20°C until MS analysis.

#### MS analysis

The vast majority of MS samples were analyzed on an EASY-nLC 1200 system (Thermo) coupled to either a Q Exactive™ HF-X Hybrid Quadrupole-Orbitrap™ mass spectrometer (Thermo) or an Orbitrap Exploris™ 480 mass spectrometer (Thermo), respectively referred to as “HF-X” and “Exploris” hereafter. The exact hardware used for each MS raw data file is defined in the experimental design template available on ProteomeXchange (PXD034888). For each run, 5 μL of sample was injected. Separation of peptides was performed using 20-cm columns (75 μm internal diameter) packed in-house with ReproSil-Pur 120 C18-AQ 1.9 µm beads (Dr. Maisch). Elution of peptides from the column was achieved using a gradient ranging from buffer A (0.1% formic acid) to buffer B (80% acetonitrile in 0.1% formic acid), at a flow of 250 nl/min. For HF-X runs, the gradient length was 100 min per sample, including ramp-up and wash-out, with an analytical gradient of 75 min ranging from 7 % B to 38 % B. For Exploris runs, the gradient length was 80 min per sample, including ramp-up and wash-out, with an analytical gradient of 57 min ranging from 7 % B to 30-36 % B depending on sample type (see experimental design template). Analytical columns were heated to 40°C using a column oven, and ionization was achieved using a Nanospray Flex Ion Source (Thermo) on the HF-X or a NanoSpray Flex™ NG ion source on the Exploris. Spray voltage set to 2 kV, ion transfer tube temperature to 275°C, and RF funnel level to 40%. Full scan range was set to 300-1,500 *m*/*z* (HF-X) or 300-1,300 *m*/*z* (Exploris), MS1 resolution to 120,000, MS1 AGC target to 3,000,000 charges (HF-X) or 2,000,000 charges (Exploris), and MS1 maximum injection time to 120 ms (HF-X) or “Auto” (Exploris). Precursors with charges 2-6 were selected for fragmentation using an isolation width of 1.3 *m*/*z*, and fragmented using higher-energy collision disassociation (HCD) with normalized collision energy of 25. Precursors were excluded from re-sequencing by setting a dynamic exclusion of 100 s (HF-X) or 80 s (Exploris). MS2 resolution was set to 45,000, MS2 AGC target to 200,000 charges, minimum MS2 AGC target to 20,000 (HF-X) or intensity threshold to 230,000 charges per second (Exploris), MS2 maximum injection time to 90 ms (HF-X) or “Auto” (Exploris), and TopN to 9. Exceptions to MS2-specific parameters are listed in the experimental design template.

#### LC-MS analysis of histone modifications

Peptides were re-suspended in 17 μl of 0.1% TFA. A total of 5.0 μl were injected into a nano-HPLC device (Thermo Fisher Scientific) using a gradient from 4% B to 90% B (solvent A 0.1% FA in water, solvent B 80% ACN, 0.1% FA in water) over 90 min at a flow rate of 300 nl/min in a C18 UHPCL column (Thermo Fisher Scientific). Data was acquired in PRM positive mode using a Q Exactive HF spectrometer (Thermo Fisher Scientific) to identify and quantify specific N-terminal peptides of histone H3 and histone H4 proteins and their PTMs. MS1 spectra were acquired in the m/z range 250-1600 with resolution 60,000 at m/z 400 (AGC target of 3×10^6^). MS2 spectra were acquired with resolution 15,000 to a target value of 2×10^5^, maximum IT 60ms, isolation 2 window 0.7 m/z and fragmented at 27% normalized collision energy. Typical mass spectrometric conditions were: spray voltage, 1.5kV; no sheath and auxiliary gas flow; heated capillary temperature, 250°C.

### Quantification and statistical analysis

#### Analysis of MS data

Triple SILAC chaperones IPs and histones IPs MS RAW data were analyzed using the version v1.6.3.4. Distinct experiments were analyzed in separate computational runs, as defined in the experimental design template available on ProteomeXchange (PXD034888). The human FASTA database used in this study was downloaded from Uniprot on the 13^th^ of May, 2019. Default MaxQuant settings were used, with exceptions specified below. Label-free quantification was enabled for all sample. Matching between runs and second peptide search were enabled. For triple SILAC samples, the re-quantify option was activated and multiplicity was set to 3, with SILAC labels set to Arg0;Lys0 (light) and Arg6;Lys4 (medium) and Arg10;Lys8 (heavy), respectively.

#### MS data analysis and Quantification of histone modifications

Raw files were searched with the Skyline software version 21.1 (Pino et al., 2020) against histone H3 and H4 peptides and their respective PTMs with a precursor mass tolerance of 5 ppm. The chromatogram boundaries of +1, +2, +3 and +4 charged peaks were validated and the Total Area MS1 under the first 4 isotopomers was extracted and used for relative quantification and comparison between experimental groups. The Total Area MS1 of co-eluting isobaric peptides (i.e., H3K36me3 and H3K27me2K36me1) was resolved using their unique MS2 fragment ions. Relative abundances (percentages) were calculated as in the following example for H3K18 acetylation:

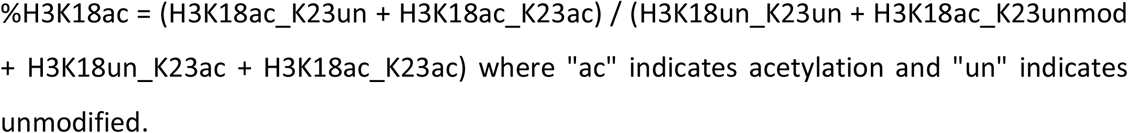

The mass spectrometry proteomics data have been deposited to the ProteomeXchange Consortium via the PRIDE partner repository with the dataset identifier PXD034924.

#### Statistical analysis of MS data

For triple SILAC chaperones IPs, MS RAW MaxQuant outputs (proteinGroups.txt) were analyzed using the Perseus software, version 1.6.14.0. For all datasets, the proteomics data was filtered to exclude potential contaminants hits, reverse-database hits, and proteins identified via modified peptides only. For all datasets, except the triple SILAC ASF1a vs ASF1b vs control, the light, medium, and heavy SILAC channels were subjected to the LFQ algorithm to accurately quantify and normalize protein ratios as derived from the ratios of individual peptides (Cox et al., 2014). In proteinGroups.txt, these values are written as LFQ intensity L, M or H. While LFQ is an acronym for Label-Free Quantification, the algorithm also accurately normalizes the individual SILAC channels, respecting both inter- and intra-experiment values derived from the different labels. The LFQ-normalized SILAC channel values were log2-transformed and filtered for detection at n=4 in at least one experimental condition. Missing values were then imputed using the default Perseus setting (down shift of 1.8 and a width of 0.3). Student’s two-sample t testing was performed with permutation-based FDR control, with s0 and FDR values stated in **Table S1** sheet for each experiment. For the triple SILAC ASF1a vs ASF1b vs control dataset, the LFQ-normalized SILAC channel values were log2-transformed and filtered for detection at n=2 in at least one experimental condition. See Table S1.

H3.1 and H3.3 datasets were analysed using label-free mass spectrometry and data was processed, filtered and Log_2_ transformed similarly to triple SILAC datasets. The resultant matrix was then split into experiment type (H3.1 and H3.3), filtered for n=5/5 valid values in at least one siRNA or siCTRL condition, median-normalized and imputed. T-tests were then performed (S0=0.1, FDR=0.05) prior to merging the H3.1 and H3.3 matrices. The MaxQuant peptide quantification output (peptides.txt) was similarly processed to extract histone peptide LFQ intensities on which T tests were also performed (S0=0.1, FDR=0.05). See **Table S1**.

For histone post-translational modifications, Total Area MS1 values were corrected for technical variability based on the abundances of the SIL heavy standard peptides across the samples. SIL standards were spiked in each sample at the same concentration, therefore any variability observed on these heavy peptides must come from technical sources. The resulting heavy-normalized intensities of the endogenous PTMs were used to calculate relative abundances by grouping PTMs that occur on the same peptide sequence. Percentages were then compared between experimental groups using unpaired two-sided t-tests. See Table S2.

#### Data visualization and network analysis

Scatter and bar plot were visualized in GraphPad Prism, version 9.0. Bubble plots and heatmaps were visualized with R studio using the libraries ggplot2 version v3.3.3, scales version 1.1.1, RColorBrewer version 1.1.2, and pheatmap version 1.0.12. Network analysis was performed in Cytoscape version 3.9.1, with the stringApp version 1.7.0, the Omics Visualizer app version 1.3.0, ClusterMaker2 app version 2.0. Venn diagrams were generated in Cytoscape with the Venn and Euler Diagrams app version 1.0.3. Functionally associated histone independent interactors (STRING score>0.6) and histone dependent (string score>0.7) were clustered using Markov Clustering (MCL, granularity=3.5) in Cytoscape using the stringApp and ClusterMaker2. Ribosomal proteins, which are common contaminants in histones purifications were excluded from network visualizations for clarity, but are listed in **Table S1**. All other statistical analysis was performed in GraphPad Prism version 9.0 and test details and p values are referred to in Figure legends.

#### Data visualization and network analysis

##### Protein complex structure predictions

Structural predictions of SPT2-H3.1–H4–ASF1a were performed using AlphaFold v2.0 (Evans *et al.*, 2021; Jumper *et al.*, 2021) in multimer mode with a maximum template date of 2021-11-01 and an input FASTA file of full-length protein sequences (UniProt IDs: P68431, P62805, Q9Y294 and Q68D10). The top five ranked models were similar in respect to the way SPT2 and ASF1 associated with H3.1–H4, and structural analysis of the highest confidence prediction, including PAE domain clustering analysis was performed using UCSF ChimeraX v1.4 (2022-04-07) (Pettersen et al., 2021).

## SUPPLEMENTAL FIGURE TITLES AND LEGENDS

**Figure S1.**
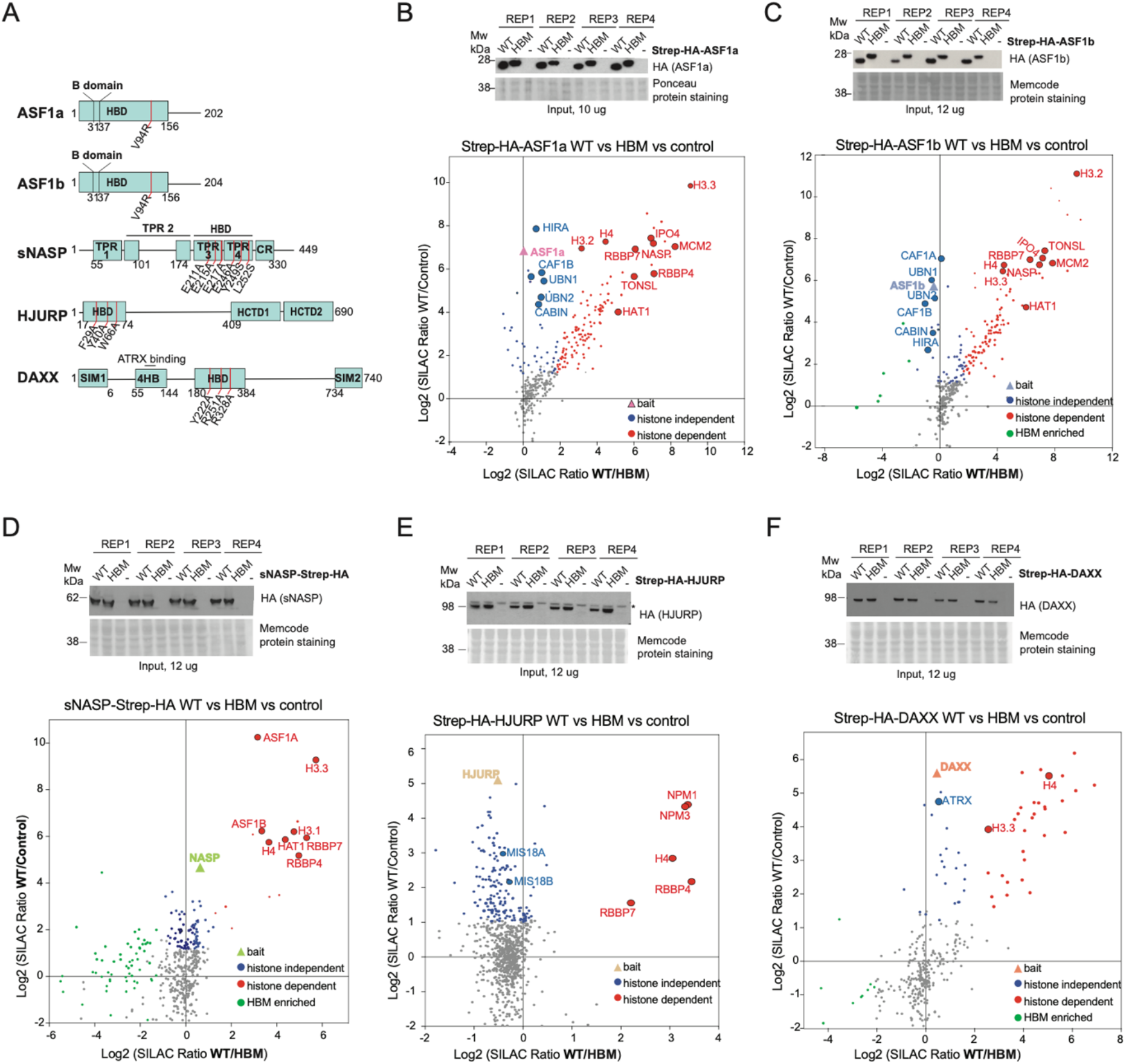
Input and IP-MS analysis of histone chaperones pulldowns – Related to Figures 1–2. (A) Schematic domain architectures of ASF1a, ASF1b, sNASP, HJURP and DAXX. Red lines indicate the point mutations in the histone binding domain (HBD) used to disrupt the protein ability to bind histone H3–H4. (B-F) Top panels show western blot analysis of soluble extract from cells expressing STREP-HA-tagged ASF1a, ASF1b, sNASP, HJURP and DAXX (WT or HBM) and control cells (−), used to perform the triple SILAC IP/MS pulldowns shown in **Figs. 1–2**. The panel show n=4 biological replicates. *, unspecific band. Bottom panels show mass spectrometry analysis of SILAC labelled pull-downs of ASF1a, ASF1b, sNASP, HJURP and DAXX, WT and HBM and control. Each pulldown was performed from soluble cell extracts, n=4 biological replicates. Proteins referred to by human Uniprot protein canonical name. The proteins nodes and names are colored according to the threshold indicated in **Table S1.** Red, blue, and green indicate histone dependent, independent, and HBM enriched factors, respectively.

**Figure S2.**
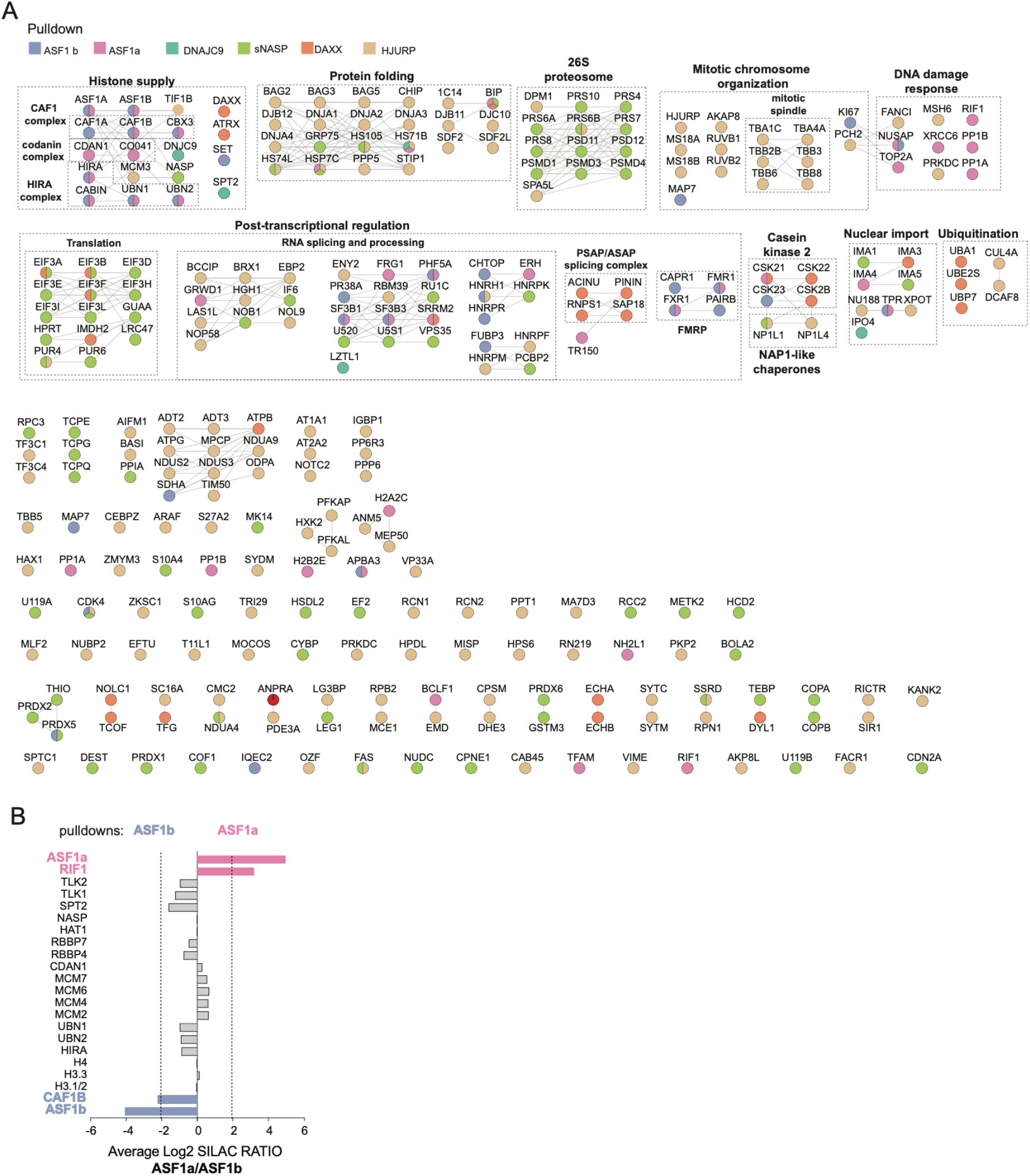
Analysis of histone independent chaperone interactome – Related to Figure 1B-C. (A) Complete list of histone independent proteins clusters as shown in Figure 1B, generated using the STRING database and MCL clustering function. Well established protein complexes and pathways are indicated by dashed black boxes and named in bold, black text. Edges indicates Protein-protein interactions according to the string database. The BAITs are indicated in bold, blue text. See **Figure S2A**. (B) Mass spectrometry analysis of SILAC labelled pull-downs of wild type ASF1a and ASF1b and control cell from soluble cell extract; n=2 biological replicates. Bar plots represent the average SILAC ratio.

**Figure S3.**
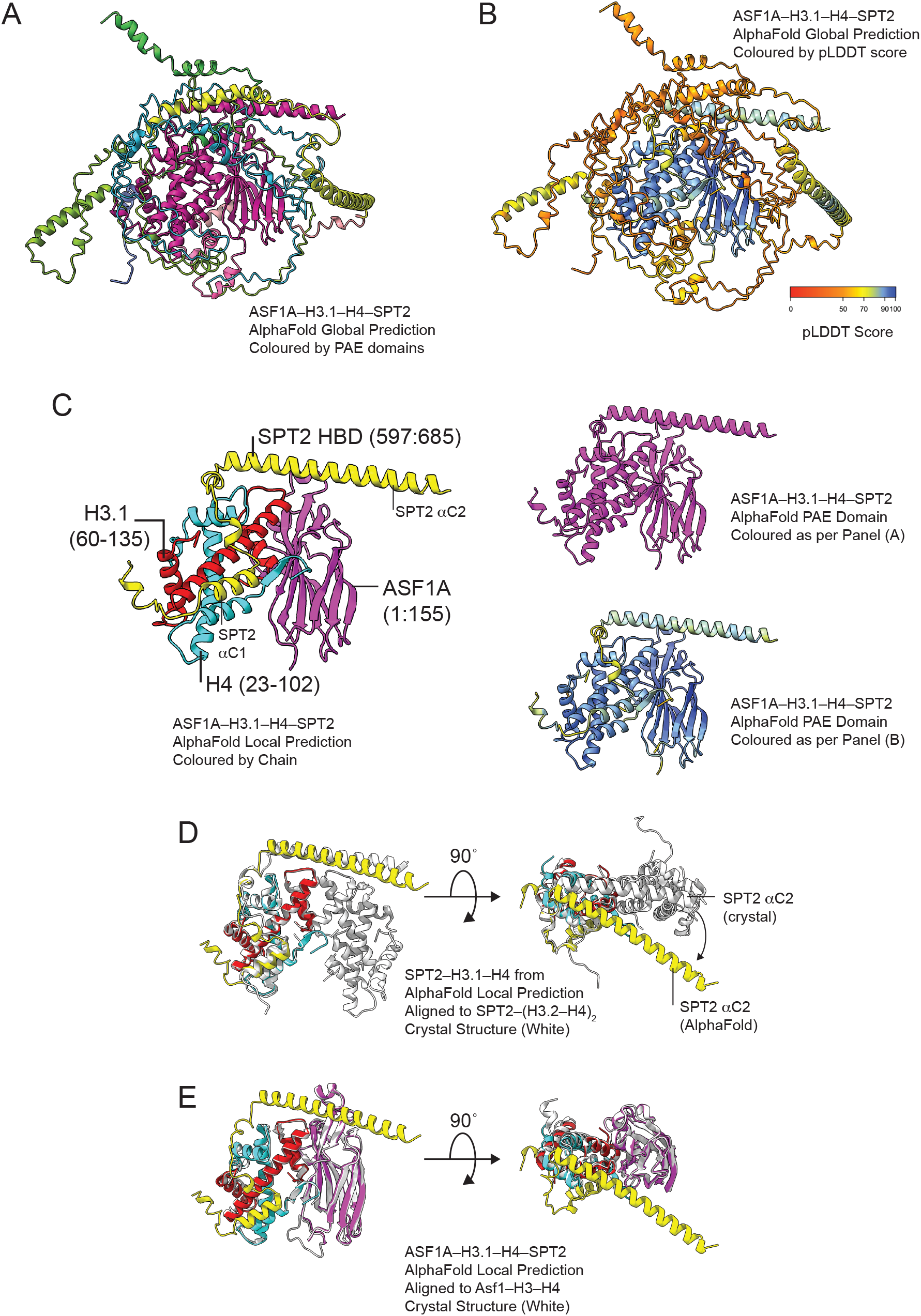
Structural characterization of ASF1-SPT2 co-chaperone complex – related to Figure 3C-D. A) ‘Global’ AlphaFold prediction of the SPT2–H3.1–H4–ASF1A histone co-chaperone complex colored by PAE domains, in magenta is the region of SPT2–H3.1–H4–SPT2 related to Figure 3C. B) ‘Global’ AlphaFold prediction of the SPT2–H3.1–H4–ASF1A histone co-chaperone complex colored by the per residue confidence score (pLLDT) showing the accuracy of locally predicted structural elements in the model. C) ‘Local’ AlphaFold prediction of SPT2 and ASF1A histone binding domains bound to H3.1–H4 extracted from full-length alpha fold prediction, colored by Left: protein chain (SPT2: yellow; ASF1A: magenta; H3.1: red; H4; blue); Top right: PAE domain; Bottom right: pLLDT score. Related to main **Figure 3C**. D) Alignment of local AlphaFold prediction of SPT2–H3.1–H4–ASF1A (colored as per panel C, with ASF1A omitted for clarity) to the crystal structure of SPT2–(H3.2–H4)2 (white; PDB: 5BS7) E) Alignment of ‘local’ AlphaFold prediction of SPT2–H3.1–H4–ASF1A (colored as per panel C) to the crystal structure of Asf1– H3–H4 (white; PDB: 2HUE) F) Alignment of ‘local’ AlphaFold prediction of SPT2–H3.1–H4–ASF1A (colored as per panel C) to the crystal structure of SPT2– (H3.2–H4)2 (white; PDB: 5BS7, with H3.2–H4 omitted for clarity) demonstrating steric clashes between ASF1A (magenta) and the SPT2 aC2 helix (white) that are released by a relocalisation of the SPT2 aC2 helix in the AlphaFold prediction (yellow).

**Figure S4.**
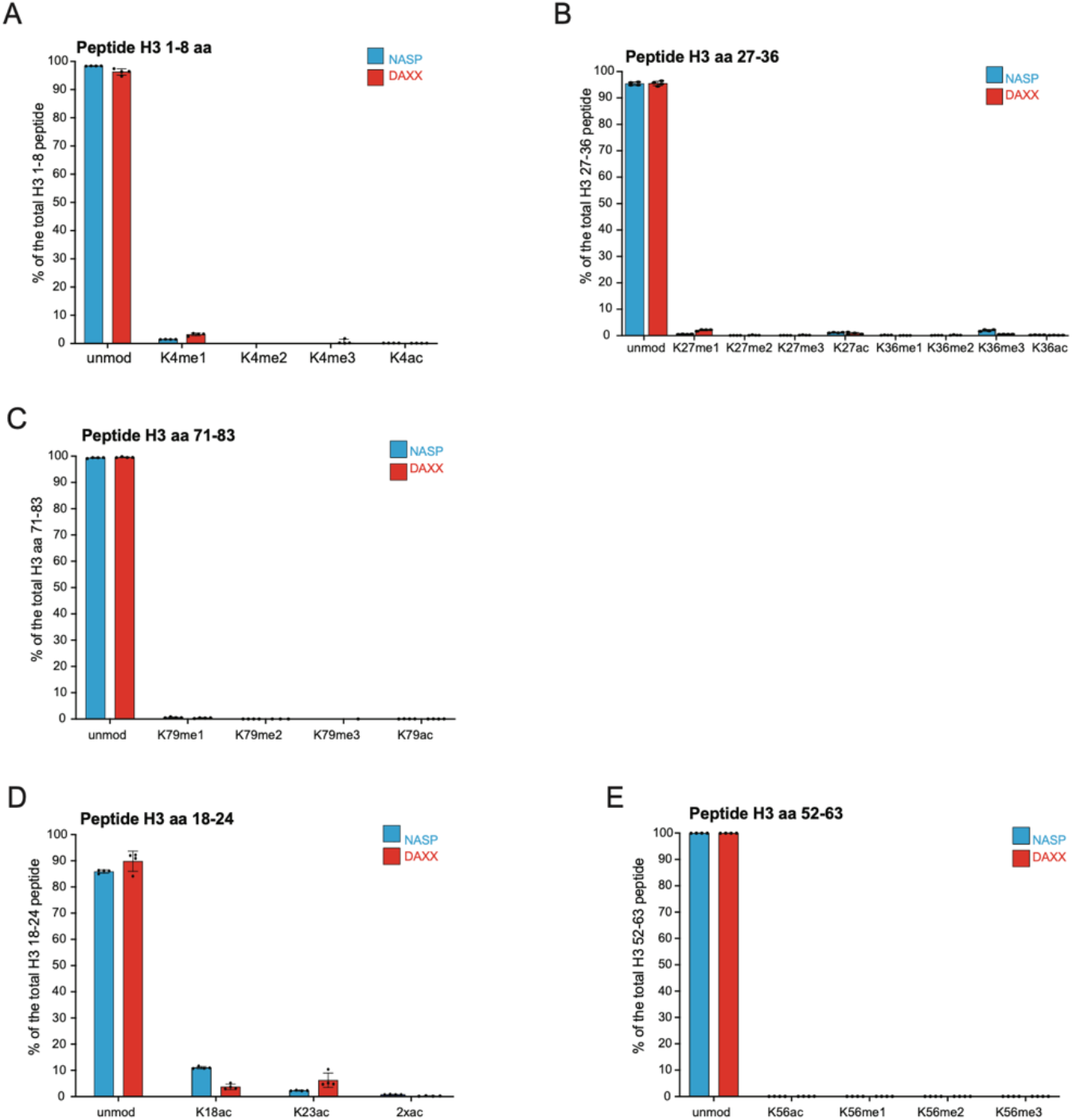
Additional profiling of marks on histones H3–H4 in DAXX and sNASP complexes – Related to Figure 5. (A-E) Analysis by quantitative mass spectrometry of histone modifications as in **Figure 6**. The graphs show averages of four biological replicates with error bars indicating SD. For additional details, see also **Table S2**. (A) Quantification of modifications on H3 peptides 1-8. (B) Quantification of modifications on H3 peptides 27-36. (C) Quantification of modifications on H3 peptides 71-83. (D) Quantification of modifications on H3 peptides 18-24. (E) Quantification of modifications on H3 peptides 52-63.

**Figure S5.**
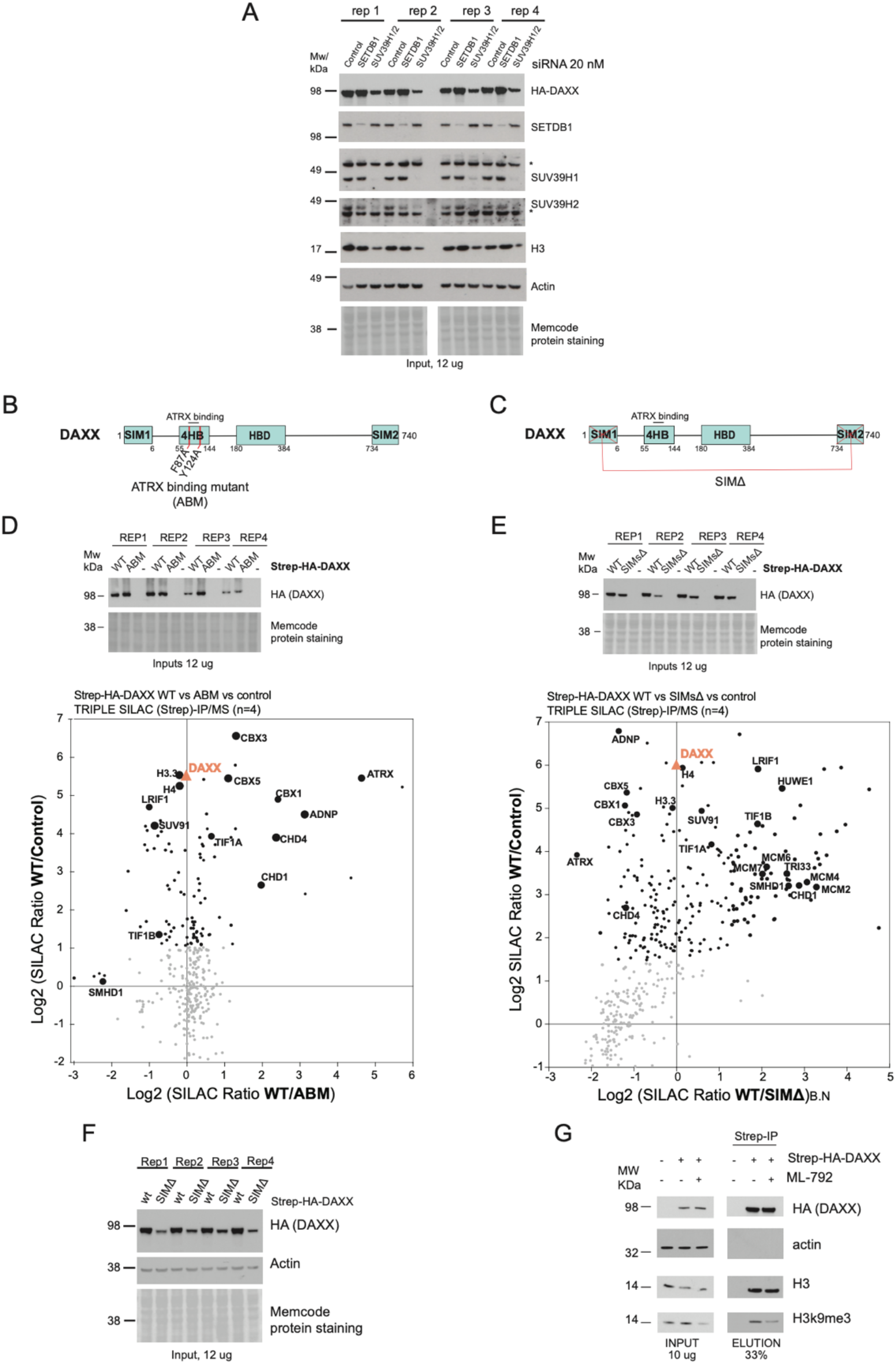
Inputs and scatter plot of DAXX ABM and SIMsΔ triple SILAC IP-MS and histone PTMs experiments – Related to Figure 6. (A) Western blot analysis of soluble extract from cells expressing Strep-HA-tagged DAXX WT and siRNA depleted for either SEDTB1 or SUV39H1/2 and compared with Control siRNA, used to perform the histone profiling shown in Figure 6A. The figure is representative of n=4 biological replicates for all the conditions. However, the control sample from the second replicate was removed from the MS analysis due to insufficient pulldown material. (B-C) Schematic domain architectures of DAXX ATRX binding mutant (ABM) and of DAXX SIMΔ mutant. Red lines indicate the point mutations in the 4-helix bundle (4HB) and the deletion of the Sumo interacting motifs 1/2 (SIM1/2), used to disrupt DAXX ability to bind ATRX. (D) Top panels show Western Blot analysis of soluble extract from cells expressing STREP-HA-tagged DAXX (WT or ABM) and control cells (−), used to perform the triple SILAC IP/MS pulldowns shown in **Figure 6C**. The panel show n=4 biological replicates. *, unspecific band. Bottom panels show mass spectrometry analysis of SILAC labelled pull-downs of DAXX, WT and ABM and control. Each pulldown was performed from soluble cell extracts, n=4 biological replicates. Proteins referred to by human Uniprot protein canonical name. The used thresholds are indicated in **Table S1.** Black indicates proteins enriched over control. (E) Top panels show Western Blot analysis of soluble extract from cells expressing STREP-HA-tagged DAXX (WT or SIMΔ) and control cells (−), used to perform the triple SILAC IP/MS pulldowns shown in **Figure 6C**. The panel show n=4 biological replicates. *, unspecific band. Bottom panels show mass spectrometry analysis of SILAC labelled pull-downs of DAXX, WT and SIMsΔ and control. The experiment was performed as in Figure **S5D.** The used thresholds are indicated in **Table S1.** Black indicates proteins enriched over control. (F) Western blot analysis of soluble extract from cells expressing Strep-HA-tagged DAXX WT or SIMΔ, used to perform the histone PTMs profiling shown in **Figure 6D**. The figure shows n=4 biological replicates. (G) Pulldown of Strep-HA-tagged DAXX from soluble fraction HeLa S3 cells induced to express either DAXX WT (+) or uninduced control cells (−). Cells were co-treated with ML-792 for 6 hours as indicated. The figure is a representative from n=2 biological replicate.

## Excel table title and legends

Table S1. Statistically processed mass spectrometry data set, related to Figure 1B,1C, 2A-B, 3A, 4A-C, 6A, S1B-F, S2A-B, S5B, S5D.

Table S2. Processed histone PTMs identified by mass spectrometry, related to Figure 5B-E, 6A-E, S4C-G.

## Notes

### Competing Interest Statement

The authors have declared no competing interest.

